# Human GBP1 differentially targets *Salmonella* and *Toxoplasma* to license recognition of microbial ligands and caspase-mediated death

**DOI:** 10.1101/792804

**Authors:** Daniel Fisch, Barbara Clough, Marie-Charlotte Domart, Vesela Encheva, Hironori Bando, Ambrosius P Snijders, Lucy M Collinson, Masahiro Yamamoto, Avinash R Shenoy, Eva-Maria Frickel

## Abstract

Guanylate binding proteins (GBPs), a family of interferon (IFN)-inducible GTPases, can promote cell-intrinsic defense by removal of intracellular microbial replicative niches through host cell death. GBPs target pathogen-containing vacuoles or the pathogen itself, and assist in membrane-disruption and release of microbial molecules that trigger cell death by activating the inflammasomes. We previously showed that GBP1 mediates atypical apoptosis or pyroptosis of human macrophages infected with *Toxoplasma gondii* (*Tg*) or *Salmonella enterica* Typhimurium (STm), respectively. In mice, the p47 Immunity-related GTPases (IRGs) control the recruitment of GBPs to microbe-containing vacuoles and subsequent cell death. However, humans are devoid of functional IRGs, and the pathogen-proximal immune detection mechanisms by GBP1 are poorly understood. Here, we describe two novel single-cell assays which show that GBP1 promotes the lysis of *Tg*-containing vacuoles and *Tg* plasma membrane, resulting in the cytosolic detection of *Tg*-DNA. In contrast, we show GBP1 only targets cytosolic STm and does not contribute to bacterial escape into the cytosol of human macrophages. GBP1 interacts with caspase-4 and recruits it directly to the bacterial surface, where caspase-4 can be activated by LPS. During STm infection, caspase-1 cleaves and inactivates GBP1 at Asp192, a site conserved in related mammalian GBP1 proteins but not in murine Gbps. STm-infected human macrophages expressing a cleavage-deficient GBP1 mutant exhibit higher pyroptosis due to the absence of caspase-1-mediated feedback inhibition of the GBP1-caspase-4 pathway. Our comparative studies elucidate microbe-specific spatiotemporal roles of GBP1 in detecting infection and the assembly and regulation of divergent caspase signaling platforms.

## Introduction

Most nucleated cells can defend themselves against infection by viruses, bacteria and eukaryotic parasites in a process called cell-intrinsic immunity. These defense programs are set in motion in response to the detection of pathogens by membrane-bound or cytosolic pattern recognition receptors (PRRs) (MacMicking, 2012; Randow et al, 2013; Mostowy & Shenoy, 2015; Jorgensen et al, 2017). In addition to antimicrobial molecules that restrict or kill pathogens, host cell-death is a destructive yet effective mechanism of defense because it removes replicative niches and traps intracellular pathogens within the cell remnants (Jorgensen et al, 2016). Antimicrobial immunity and cell death are enhanced by the type II interferon (IFNγ) which induces the expression of up to 2000 IFN-stimulated genes (ISGs) (MacMicking, 2012; Schoggins, 2019). The guanylate binding protein (GBP) family of GTPases, which are highly abundant in cells exposed to IFNγ, consists of seven members in the human and eleven members in mice (Olszewski et al, 2006; Shenoy et al, 2007; Kresse et al, 2008; Shenoy et al, 2012). GBPs target intracellular pathogens and mediate host-defense through multiple mechanisms, including the regulation of autophagy, oxidative responses, inflammasomes and cell death (Jorgensen *et al*, 2016; Olszewski *et al*, 2006; Tripal *et al*, 2007; Kresse *et al*, 2008; Kim *et al*, 2011, 2012b; MacMicking, 2012; Shenoy *et al*, 2012; Haldar *et al*, 2013; Randow *et al*, 2013; Haldar *et al*, 2014; Meunier *et al*, 2014; Haldar *et al*, 2015; Man *et al*, 2015; Meunier *et al*, 2015; Mostowy & Shenoy, 2015; Shenoy *et al*, 2007; Feeley *et al*, 2017; Foltz *et al*, 2017; Jorgensen *et al*, 2017; Li *et al*, 2017; Lindenberg *et al*, 2017; Man *et al*, 2017; Piro *et al*, 2017; Wallet *et al*, 2017; Wandel *et al*, 2017; Zwack *et al*, 2017; Costa Franco *et al*, 2018; Liu *et al*, 2018; Santos *et al*, 2018; Gomes *et al*, 2019; Schoggins, 2019).

Once GBPs target to a pathogen vacuole or the pathogen itself, they are thought to disrupt these membranes by an as yet uncharacterized mechanism (Yamamoto et al, 2012; Selleck et al, 2013; Meunier et al, 2014; Kravets et al, 2016). Disruption of barrier membranes leads to pathogen growth-control and release of pathogen-associated molecular patterns (PAMPs) which are sensed by PRRs that can trigger host cell death. Whether GBPs directly recognize pathogen vacuolar membranes or PAMPs is an important question that has not yet been answered (Pilla et al, 2014; Meunier et al, 2014; Lagrange et al, 2018; Santos et al, 2018; Fisch et al, 2019a).

A large body of work on GBPs has been carried out in murine cells, wherein these proteins closely collaborate with members of a second family of IFN-induced GTPases, comprising 23 members in the mouse, the p47 immunity-related GTPases (IRGs) (MacMicking et al, 2003; Bernstein-Hanley et al, 2006; Singh et al, 2006; Henry et al, 2007; Miyairi et al, 2007; Shenoy et al, 2007; Coers et al, 2008; Hunn et al, 2008; Al-Zeer et al, 2009; Tiwari et al, 2009; Khaminets et al, 2010; Lapaquette et al, 2010; Singh et al, 2010; Brest et al, 2011; Haldar et al, 2014). For instance, mouse IrgB10 targets bacteria following mGbp recruitment and contributes to the release of bacterial LPS and DNA, and mouse IrgM1 and −3 are essential regulators of GBP-targeting of some pathogen-containing vacuoles (Singh et al, 2010; Meunier et al, 2014; Haldar et al, 2015; Man et al, 2016; Balakrishnan et al, 2018). However, only one IFN-insensible, truncated IRG, IRGM, can be found in the human genome (Bekpen et al, 2005, 2010). Therefore, how human GBPs target intracellular pathogens remains unknown. In addition, some PRRs are unique to humans, for example, LPS-sensing by both caspase-4 and caspase-5 in humans but only caspase-11 in the mouse (Kayagaki et al, 2011, 2013; Hagar et al, 2013; Shi et al, 2014; Casson et al, 2015; Ding & Shao, 2017). Moreover, unlike mouse cells, human cells can respond to tetra-acylated LPS (Lagrange et al, 2018) and possess additional DNA sensors, such as the DNA-dependent protein kinase (Burleigh et al, 2020). The mechanisms underlying GBP-mediated detection of pathogens and stimulation of human macrophage death therefore need to be investigated further.

All seven human GBPs have a conserved structure with an N-terminal globular GTPase domain and a C-terminal helical domain. GBP1, GBP2 and GBP5 are isoprenylated at their C-terminal CaaX-box, which can anchor them to membranes (Nantais et al, 1996; Olszewski et al, 2006; Tripal et al, 2007; Britzen-Laurent et al, 2010). The ability of human GBPs to target pathogen-containing vacuoles remains poorly characterized. Differences have also been noted depending on the pathogen and cell type. We previously showed that human GBP1 fails to target the apicomplexan parasite *Toxoplasma gondii* (*Tg*) and two intracellular bacterial pathogens, *Chlamydia* and *Salmonella enterica* subsp. *enterica* serovar Typhimurium (STm), in human A549 epithelial cells; however, GBP1 is required for the restriction of parasite growth, but not the bacterial pathogens (Johnston et al, 2016). On the other hand, in human macrophages GBP1 localizes to *Tg*, *Chlamydia* and STm, but whether it can disrupt membranes that enclose these pathogens is not known (Al-zeer et al, 2013; Fisch et al, 2019a).

Human GBP1 targets *Tg* and STm and promotes distinct forms of macrophage cell death. In the case of both pathogens, GBP1 targeting to pathogens is necessary, even though downstream mechanisms of cell death are distinct. Since *Tg* induces the loss of inflammasome proteins, including NLRP3 and caspase-1, human macrophages undergo atypical apoptosis through the assembly of AIM2-ASC-caspase-8 complexes. In contrast, GBP1 promotes activation of caspase-4 following its recruitment to STm resulting in enhanced pyroptosis (Fisch et al, 2019a). Although our previous work suggested that GBP1 is involved in PAMP release for detection by these PRRs during natural infection (Fisch et al, 2019a), the underlying mechanisms involved in liberating microbial ligands was not investigated.

In this study we show that GBP1 contributes to lyses of the parasite-containing vacuole and the plasma membrane of *Tg* by employing two newly developed assays. We also show that GBP1 only targets STm that are already cytosolic and does not contribute to their ability to reach the cytosol of human macrophages. In contrast, during STm infection, caspase-1 cleaves and inactivates GBP1 and thereby reduces its ability to recruit caspase-4. These studies reveal the feedback inhibition of GBP1/caspase-4-driven pyroptosis during STm infection and its dual membrane-disruptive actions during *Tg* infection.

## Results

### GBP1 contributes to *Toxoplasma* parasite and vacuole disruption and infection control

As GBP1 elicits divergent host cell death programs in response to *Tg* and STm, we sought to investigate the upstream mechanistic details of GBP1 during infection by these two unrelated human pathogens. We previously correlated GBP1 recruitment to *Tg* parasitophorous vacuoles (PV) to activation of AIM2 and caspase-8 with the recognition of parasite DNA (Fisch et al, 2019a). Like some murine Gbps (Yamamoto et al, 2012; Selleck et al, 2013; Degrandi et al, 2013; Kravets et al, 2016), we therefore hypothesize that human GBP1 promotes PV opening and cytosolic access to intravacuolar pathogens.

Extending our previous finding of GBP1 recruiting to the PV, we also localized GBP1 directly to the surface of *Tg* using AiryScan super-resolution microscopy (**Figure 1A**). To test whether GBP1 can disrupt *Tg* PVs and potentially the parasites, we used cytosolic dye CellMask, which is excluded from PVs but enters once the PV membrane (PVM) is disrupted (**Figure 1B**). As positive control for this new assay, vacuoles were chemically disrupted by detergent-mediated permeabilization resulting in higher fluorescence within the vacuoles as compared to untreated cells (**Figure 1B**). Increased CellMask dye intensity within naturally disrupted *Tg* vacuoles could be reliably quantified using our artificial intelligence-based high-throughput image analysis workflow HRMAn (Fisch et al, 2019b), which enabled us to enumerate dye access within thousands of PVs formed upon infection of type I and type II strains of *Tg.* Analyses of CellMask fluorescence within PVs in IFNγ-primed THP-1 wildtype (WT) cells revealed increased intensities, indicating their disruption (**Figure S1**). IFNγ-primed THP-1 Δ*GBP1* cells showed that *Tg* vacuoles were not disrupted, as seen by the exclusion of CellMask dye (**Figure S1**). Doxycycline induced reexpression of GBP1 (THP-1 Δ*GBP1*+Tet-*GBP1* cells) rescued vacuole breakage; as control, empty vector transduced cells (THP-1 Δ*GBP1*+Tet-EV) behaved like Δ*GBP1* cells (**Figure S1**). We next used Doxycycline-induced expression of mCherry-GBP1 (THP-1 Δ*GBP1*+Tet-mCH-*GBP1* cells) to allow quantification of GBP1-recruitment to *Tg* and stratify data on whether PVs that were decorated with mCH-GBP1 lost their integrity. Indeed, a population of GBP1^+^ PVs were unable to exclude CellMask dye clearly indicating loss of membrane integrity (**Figure 1C**). Taken together, we conclude that GBP1 is contributing to opening of PVs and GBP1-targeted vacuoles preferentially undergo loss of membrane integrity.

**Figure 1:**
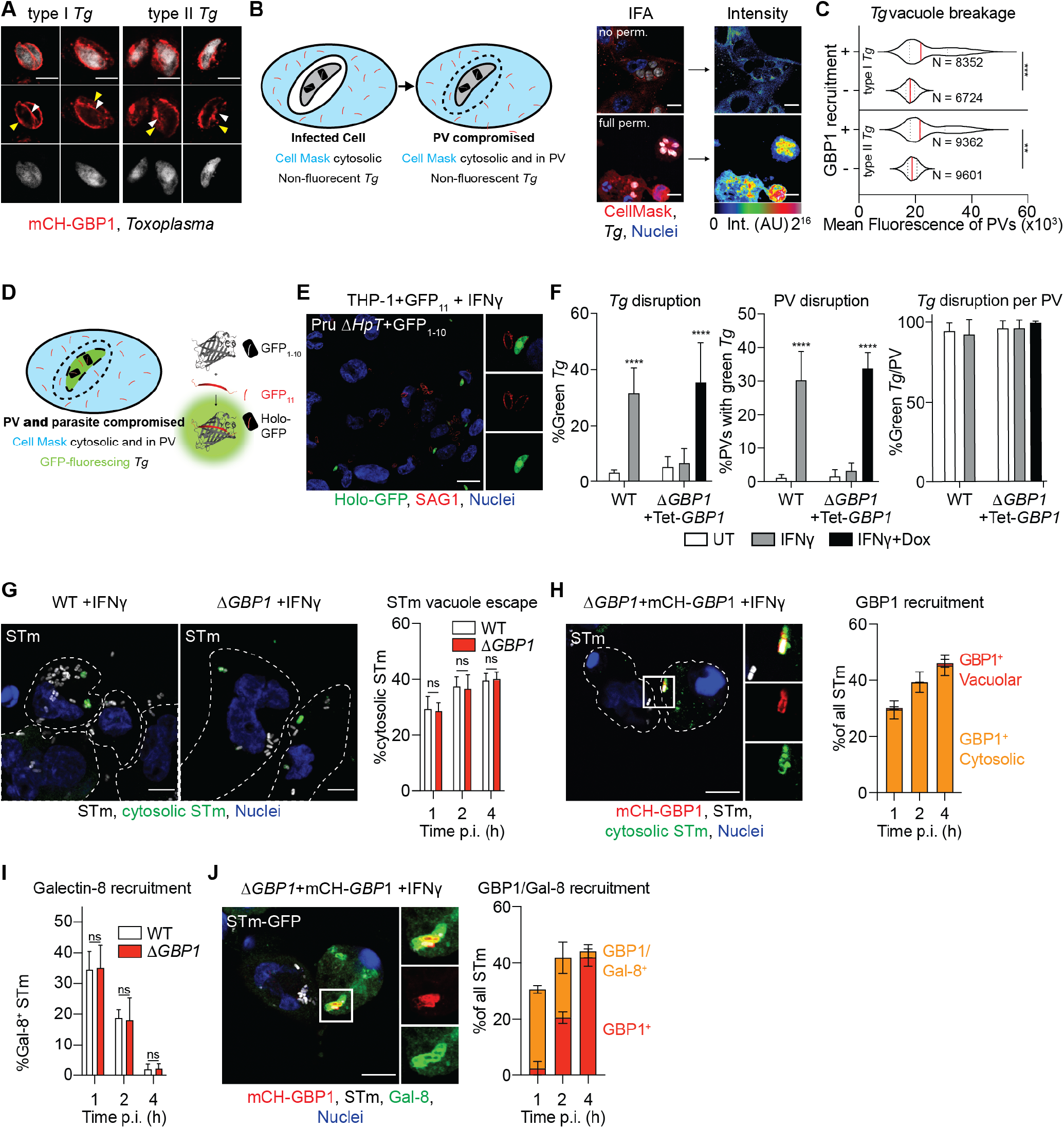
GBP1 disrupts Toxoplasma vacuoles and parasite membrane, and targets cytosolic *Salmonella*. **(A)** AiryScan immunofluorescence images of type I or type II *Toxoplasma gondii* (*Tg*) decorated with mCH-GBP1 in IFNγ and Doxycycline (Dox) treated THP-1 Δ*GBP1*+Tet-mCH-*GBP1* cells. Red: mCH-*GBP1*; White: *Tg*. White arrowhead indicates GBP1 on the parasite and yellow arrowhead indicates GBP1 on the vacuole membrane. Scale bar 4 μm. **(B)** Left: Illustration of novel high-throughput imaging assay to measure parasitophorous vacuole (PV) integrity by Cell Mask flooding and Right: Representative immunofluorescence images from proof-of-principle experiment using THP-1 WT infected with type I *Tg* for 18 hours and stained with CellMask but not permeabilized (no perm.; Top) or fully permeabilized with saponin (full perm.; Bottom) and corresponding rainbow intensity diagram to illustrate the CellMask signal from images used for quantification; AU: arbitrary fluorescence intensity values. Red: CellMask; Grey: *Tg*; Blue: Nuclei. Scale bars 10 μm. **(C)** Representative quantification of CellMask fluorescence intensities within vacuoles (PV) of type I or type II *Tg* infected THP-1 Δ*GBP1*+Tet-mCH-*GBP1* pre-treated with IFNγ and Dox to induce GBP1 expression. Plotted depending on whether PVs were decorated with GBP1 (+) or not (-). N = number of vacuoles. **(D)** Illustration of novel high-throughput imaging assays to measure *Tg* integrity with the split-GFP system. **(E)** Example immunofluorescence image and **(F)** quantification of disrupted and thus green-fluorescing type II *Tg* parasites expressing GFP_1-10_ fragment (Pru Δ*Hpt*+GFP_1-10_) infecting IFNγ-primed THP-1+GFP_11_ or IFNγ and Dox-primed THP-1 Δ*GBP1*+Tet-*GBP1*+GFP_11_ cells stained for all *Tg* using anti-surface-antigen 1 (SAG1). Data plotted as proportion of all parasites (left), proportion of all PVs containing at least one green parasite (middle) or proportion green parasites within the same PV (right). Red: SAG1; Green: Holo-GFP; Blue: Nuclei. Scale bar 20 μm. **(G)** Representative immunofluorescence images at 2 hours p.i. and quantification of the proportion of cytosolic STm from differentially permeabilized, IFNγ-primed THP-1 WT or Δ*GBP1* cells infected with STm SL1344 (MOI = 30) at indicated time p.i. Cells are outlined by the white, dashed line. Grey: STm; Green: Pseudo-colored cytosolic and extracellular STm; Blue: Nuclei. Scale bar 10 μm. **(H)** Representative immunofluorescence images at 2 hours p.i and quantification of GBP1 recruitment to cytosolic and intra-vacuolar STm in IFNγ-primed and Dox-treated THP-1 Δ*GBP1*+Tet-mCH-*GBP1* infected with STm SL1344 (MOI = 30) at indicated times p.i from differentially permeabilized cells stained for cytosolic STm and total STm. Cells are outlined by the white, dashed line. Red: mCH-GBP1; Grey: STm; Green: Cytosolic STm; Blue: Nuclei. Scale bar 10 μm. **(I)** Quantification of galectin-8 (Gal-8) recruitment to STm SL1344-GFP in IFNγ treated THP-1 WT or Δ*GBP1* at the indicated times post infection. **(J)** Representative immunofluorescence images at 1 hour and quantification of Gal-8 recruitment to STm in IFNγ and Dox treated THP-1 Δ*GBP1*+Tet-mCH-*GBP1* infected with STm SL1344-GFP (MOI = 30) at the indicated time post infection. Red: mCH-GBP1; Grey: STm; Green: Gal-8; Blue: Nuclei. Scale bar 10 μm. **Data information:** Graphs in **(C)**, **(F)**, **(G)**, **(H)**, **(I)** and **(J)** show mean ± SEM from n = 3 independent experiments. ** *P* ≤ 0.01; *** *P* ≤ 0.001 or **** *P* ≤ 0.0001 in **(C)** from nested t-test comparing GBP1^+^ to GBP1^-^ PVs following adjustment for multiple comparisons, in (F) from two-way ANOVA comparing to untreated condition and in **(G)** and **(I)** from one-way ANOVA following adjustment for multiple comparisons; ns, not significant.

Elegant microscopy previously localized murine Gbps directly onto the surface of the *Tg* plasma membrane (Kravets et al, 2016) similar to our finding of GBP1 recruiting directly to the parasite surface in presumably broken vacuoles. Whether direct recruitment of a GBP to a *Tg* parasite leads to disruption of the parasite plasma membrane has not been studied. We developed a second novel assay that measures parasite membrane integrity (**Figure 1D**). In a split-GFP complementation approach, *Tg* parasites only fluoresce upon access of a GFP_11_ fragment expressed in host cell cytosol (**Figure S2A**) to the GFP_1-10_ fragment expressed in the *Tg* cytosol (**Figure S2B**); neither fragment is fluorescent on its own (Romei & Boxer, 2019). If the PV and the *Tg* membranes are both disrupted, the fragments should assemble to form fluorescent GFP holo-protein (**Figure 1D**). Indeed, we could observe GFP-fluorescing parasites in IFNγ-primed THP-1 cells (**Figure 1E**) and quantify the proportion of parasites with GFP fluorescence using high-throughput imaging (**Figure 1F**). This revealed that *Tg* only become disrupted in the presence of GBP1 (**Figure 1F**). Moreover, all parasites within the same vacuole were disrupted suggesting that once PV integrity is lost the *Tg* within them are susceptible to membrane disruption (**Figure 1F**). The disruption of parasites was further investigated using flow cytometry of *Tg* from infected THP-1 cells which showed that parasites disrupted at 6 hours post infection (p.i.) or later (**Figure S2C**). Plaque assay of sorted parasites confirmed that green-fluorescing *Tg* were not viable (**Figure S2D**).

We validated our PV disruption assays by examining the ultrastructure of the vacuole membranes using correlative light and electron microscopy, which revealed ruffled and broken vacuole membranes in cells expressing GBP1 (**Figure S3**). In THP-1 Δ*GBP1*, most PVs analyzed by electron microscopy did not show structural defects or loss of membrane integrity (**Figure S3**). Together, our novel assays indicated that GBP1 contributes to disruption of both the membrane of the PV and the plasma membrane of *Tg* parasites.

### GBP1 does not participate in *Salmonella* vacuolar escape but targets cytosolic bacteria

Having established an indispensable role for GBP1 in opening of *Tg* PVs and parasites, we wanted to test if GBP1 also contributed to the escape of STm from *Salmonella*-containing vacuoles (SCVs). In murine cells, Gbps have been found to both disrupt STm vacuoles, as well as directly recognize bacterial LPS in the cytoplasm (Meunier et al, 2014; Pilla et al, 2014). We used differential permeabilization (Meunier et al, 2014) to determine whether the escape of STm from its vacuole into the cytosol required GBP1. Similar number of cytosolic STm were detected in WT and Δ*GBP1* cells, suggesting that GBP1 is dispensable for cytosolic escape of STm (**Figure 1G**). This indicated that GBP1 has a microbe-specific role in disruption of microbial compartments. Importantly, differential permeabilization revealed that GBP1 was exclusively recruited to cytosolic STm at all time points (**Figure 1H**). Although these results in human macrophages contrast findings of mGbp involvement in STm infection of mouse macrophages (Meunier et al, 2014), they are in agreement with the lack of a role for GBPs in the opening of bacterial pathogen-containing vacuoles, for example during infection by *Legionella* (Creasey & Isberg, 2012; Pilla et al, 2014; Feeley et al, 2017; Liu et al, 2018) and *Yersinia* (Feeley et al, 2017), or directly targeting cytosolic *Francisella novicida* (Meunier et al, 2015; Man et al, 2016). In a second assay to analyze the capacity of GBP1 to open STm vacuoles, we used the lectin galectin-8 (Gal-8) as a marker for cytosolic bacteria. Studies in human epithelial cells have shown that Gal-8 is recruited to disrupted SCVs, which promotes bacterial xenophagy and growth-restriction(Thurston et al, 2012). Consistent with the previously observed lack of a role for GBP1 in cytosolic escape of STm, similar proportions of STm were decorated with Gal-8 in WT and Δ*GBP1* cells (**Figure 1I**). Temporal studies showed that SCVs were rapidly disrupted (become Gal-8^+^), but lose this marker over time (**Figure 1I**), as has been shown before in epithelial cells (Thurston et al, 2012). At later time points as the proportion of Gal-8+ vacuoles decreased, cytosolic STm still retained GBP1 coating (**Figure 1J**). These single-cell assays revealed that unlike during *Tg* infection, GBP1 does not contribute to cytosolic escape of STm, but recruits directly to cytosolic STm.

### GBP1 promotes access to PAMPs for cytosolic host defense and interacts with caspase-4 on the surface of STm

As *Tg* infection activates the DNA sensor AIM2 and we demonstrated that GBP1 promotes PV and *Tg* plasma membrane disruption, we wanted to visualize release of *Tg*-DNA into the cytoplasm of infected macrophages and subsequent recognition by AIM2. To this end, we labelled *Tg*-DNA with EdU by growing them in human foreskin fibroblasts (HFFs), whose DNA remains unlabeled as they do not replicate due to contactdependent growth inhibition. Following infection of macrophages with EdU-labelled *Tg*, we visualized *Tg*-DNA with Alexa Fluor 647 dye using click-chemistry and quantified macrophages containing cytosolic *Tg*-DNA (**Figure 2A**). Approximately 35% of infected macrophages that had at least one PV targeted by GBP1 (GBP1^+^) contained *Tg*-DNA in their cytosol at 6h p.i. while uninfected macrophages or infected macrophages without targeted PVs did not show this phenotype (**Figure 2A**). Infection of myc-AIM2 expressing THP-1 macrophages (**Figure S4C**) furthermore showed association of the cytosolic DNA sensor with EdU-labelled *Tg*-DNA in the cytosol (**Figure 2A**). Taken together, these results corroborate the model that human GBP1 actively ruptures the *Tg* PV and parasites and releases *Tg*-DNA into the cytosol for downstream detection by AIM2.

**Figure 2:**
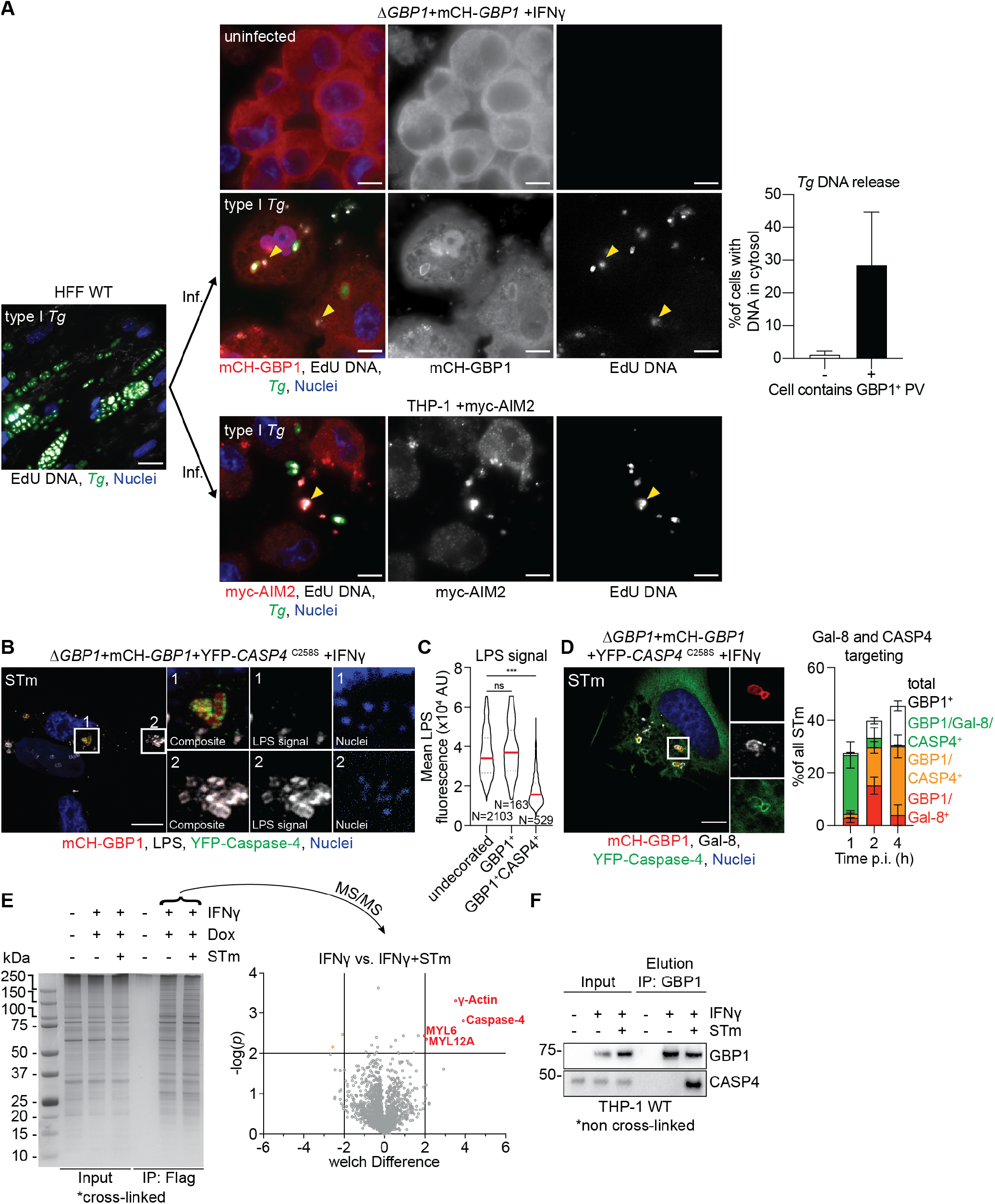
GBP1 mediates access to PAMPs during *Toxoplasma* and *Salmonella* infection. **(A)** Left: Representative immunofluorescence image of type I *Toxoplasma gondii* (*Tg*) grown in human foreskin fibroblasts (HFF WT) in the presence of EdU DNA label. Labelled type I *Tg* were harvested from the HFFs and used to infect (Inf.) THP-1 Δ*GBP1*+Tet-mCH-*GBP1* or THP-1 +*myc-AIM2* for 6 hours. THP-1 cells were pre-treated with IFNγ and Doxycycline (Dox) to induce mCH-GBP1 expression Middle: Parasite DNA released into the cytoplasm was visualized by click-chemistry to label the incorporated EdU. Right: Quantification of proportion of infected cells with cytosolic *Tg*-DNA based grouped based on whether the cells contain a GBP1-targeted (GBP1^+^) *Tg* parasitophorous vacuole (PV). Red: mCH-GBP1 or immune-stained myc-AIM2; White: EdU-DNA; Green: *Tg*; Blue: Nuclei. Some released, cytosolic (and additionally AIM2-bound, lower panels) *Tg*-DNA indicated by yellow arrowheads. Scale bar 10 μm. **(B)** Representative immunofluorescence images and **(C)** quantification of LPS staining intensity of STm from IFNγ and Dox-treated THP-1 Δ*GBP1*+Tet-mCH-*GBP1*+YFP-*CASP4*^C258S^ cells infected with STm SL1344 (MOI = 30) for 2. Red: mCH-GBP1; Grey: STm-LPS; Green: YFP-caspase-4; Blue: Nuclei. Scale bar 10 μm. Contrast enhanced in the Nuclei inset, to visualize STm-DNA used for detection of bacteria that do not stain for LPS. **(D)** Representative immunofluorescence images at 1 hour p.i. and quantification of galectin-8 (Gal-8) and caspase-4 (CASP4) recruitment to STm in IFNγ-primed and Dox-treated THP-1 Δ*GBP1*+Tet-mCH-*GBP1*+YFP-*CASP4*^C258S^ infected with STm SL1344 (MOI = 30) at indicated times p.i. Red: mCH-GBP1; Grey: Galectin-8; Green: YFP-caspase-4; Blue: Nuclei. Scale bar 10 μm. **(E)** Left: Silver stain of SDS-PAGE gel showing immunoprecipitation (IP) of Flag-GBP1 from THP-1 Δ*GBP1*+Tet-Flag-*GBP1* treated with IFNγ and Dox and infected with STm for 2 hours of left untreated. Right: Volcano plot of mass spectrometry hits comparing IFNγ treated cells with IFNγ treated and STm-infected cells. Plotted as welch difference of mass spectrometry intensities versus −log10(*P*). Significant hits shown in orange/red circles. **(F)** Representative immunoblots of immunoprecipitation of endogenous GBP1 from IFNγ-primed or naïve THP-1 WT infected with STm for 2 hours as indicated showing co-precipitation of endogenous caspase-4 identified as hit using mass spectrometry. **Data information:** Graphs in **(A)**, **(C)** and **(D)** show mean ± SEM from n = 3 independent experiments. *** *P* ≤ 0.001 in **(C)** from nested one-way ANOVA comparing to undecorated vacuoles for the means of the n = 3 independent experiments; ns, not significant.

GBP1 thus promotes the sensing of PAMPs and formation of cytosolic signaling platforms also known as supramolecular organizing centers (SMOCs) (Kagan et al, 2014). We investigated the structure of caspase activation SMOCs promoted by GBP1 actions using structured illumination microscopy (SIM). Upon *Tg* infection, AIM2 detects released *Tg*-DNA and nucleates the formation of an inflammasome containing ASC and caspase-8. Using SIM, we found that these atypical inflammasome complexes appear similar to previously described inflammasomes containing ASC and caspase-1 or caspase-8 (Man et al, 2013, 2014). We found a “donut”-like ASC ring enclosing caspase-8 in the center (**Figure S4A-B**). As we could not detect endogenous AIM2 by immunofluorescence microscopy, we resorted to using THP-1 cells expressing myc-AIM2 (**Figure S4C**), which revealed AIM2 recruitment to ASC specks in *Tg*-infected macrophages (**Figure S4D-E**). Altogether, these studies confirm that *Tg*-DNA is present within the macrophage cytosol as a result of GBP1-mediated disruption of the PVM and *Tg* membrane resulting in AIM2 activation.

We next decided to contrast GBP1 actions during STm infection, where we previously showed that caspase-4 is directly targeted to GBP1^+^ STm (Fisch et al, 2019a). The question was whether there is an interaction between GBP1 and caspase-4, which leads to recruitment of the LPS-sensor, and whether caspase-4 is recruited directly on the surface of STm. Indeed, 3D-rendered SIM imaging demonstrated GBP1 recruited caspase-4 directly to the surface of STm (**Figure S4F**). Bacteria were completely covered in GBP1, with a high degree of colocalisation with YFP-CASP4^C258S^ (**Figure S4F**). Interestingly, immunofluorescence staining of *Salmonella-LPS* using a monoclonal antibody revealed that GBP1-CASP4^+^ bacteria stained not at all or poorly, suggesting access to the epitope to be blocked (**Figure 2B+C**). STm not decorated with caspase-4 but positive for GBP1 however were stained with anti-LPS antibody (**Figure 2B**). As caspase-4 can directly bind LPS with its CARD (Shi et al, 2014), this finding is consistent with the possibility that caspase-4 on the bacterial surface precludes antibody-mediated staining of LPS (**Figure 2C**).

Caspase-4 recruitment to bacteria mirrored that of GBP1. The majority of cytosolic (Gal-8^+^) GBP1 ^+^ STm were also positive for caspase-4. Notably, GBP1-caspase-4 were retained on STm over time even though Gal-8 staining had reduced (**Figure 2D**), which suggested that GBP1-caspase-4 are present on cytosolic STm longer during infection.

Our previous work showed that the translocation of GBP1 to STm and enhanced pyroptosis requires its GTPase function and isoprenylation (Fisch et al, 2019a) but did not determine whether these effects led to deficient caspase-4 targeting to STm. THP-1 *ΔGBP1* cells reconstituted with Dox-inducible variants of GBP1 that lacked GTPase activity (GBP1^K51A^) or isoprenylation sites (GBP1^C589A^ or GBP1^Δ589-592^; **Figure S5A**) revealed that none of these variants supported the recruitment of caspase-4 (**Figure S5B+C**). Taken together, through single-cell comparative analyses we have established that GBP1-targeting to *Tg* promotes the release of parasite DNA into the cytosol, whereas GBP1-targeting to STm enables caspase-4 recruitment to cytosolic bacteria. The reduced LPS staining on bacteria further suggest that GBP1 facilitates access to bacterial LPS ligand to caspase-4.

We additionally decided to use an unbiased proteomics approach to identify GBP1 binding-partners and other proteins recruited to GBP1-caspase-4 SMOCs on cytosolic STm. For this, we immunoprecipitated Dox-inducible Flag-GBP1 from STm-infected THP-1 *ΔGBP1* cells following protein cross-linking (**Figure 2E and Figure S6**). Comparing infected to uninfected cells and correcting for nonspecific binding of proteins to the Flag-beads, identified several proteins that were enriched in infected samples above the significance cut-off (*P ≤ 0.01*, **Figure 2E**). Some of these proteins are known GBP1 interacting proteins such as γ-Actin (*ACTG1*), Myosin light polypeptide 6 (*MYL6*) and Myosin regulatory light chain 12a (*MYL12A*) (Ostler et al, 2014; Forster et al, 2014). The most prominent infection-specific GBP1 interaction partner we detected was caspase-4, which supported results from microscopy. We did not identify any other proteins that may be interacting with GBP1 and caspase-4 on the STm surface. To establish that the detected interaction is physiologically relevant during infection, we repeated immunoprecipitation experiments using antibodies against endogenous GBP1 from THP-1 WT cells (this time without prior cross-linking). In agreement with our proteomics results, endogenous GBP1 interacted with caspase-4 only during STm infection pointing towards its specific and crucial role in enabling LPS-sensing by caspase-4 (**Figure 2F**).

Taken together these results indicate that GBP1 has two modes of assembling caspase-containing complexes depending on the infecting pathogen: (1) by proxy through *Tg* vacuole and parasite membrane disruption and release of *Tg*-DNA into the cytosol to trigger activation of the AIM2 inflammasome and (2) by direct recruitment and interaction with caspase-4 on the surface of STm.

### Caspase-1, but not caspase-4, can cleave GBP1 during infection

The noncanonical inflammasome in mouse macrophages involves sequential activation of capsase-4/11 and caspase-1, wherein caspase-4/11 activation precedes caspase-1. As both caspases are independently activated in IFNγ-stimulated human macrophages infected with STm (Fisch et al, 2019a) we wanted to investigate whether a crosstalk existed between the two pathways. This was also pertinent given a previous report of caspase-1-mediated cleavage of GBP1 in human umbilical vein endothelial cells (HUVECs) (Naschberger et al, 2017); however, the functional consequences of GBP1 proteolysis during infection were not investigated in that study. Noncanonical inflammasome activation during LPS-transfection is a cell-intrinsic process that involves K^+^ efflux-mediated activation of caspase-1 (Kayagaki et al, 2011; Rühl & Broz, 2015), and release of inflammasome specks can activate inflammasomes in neighboring cells (Venegas et al, 2017). We therefore first wanted to verify that caspase-1 and caspase-4 are activated within the same STm-infected macrophage. Our results showed a perfect correlation between bacterial targeting by GBP1-CASP4 and pyroptosis and we indirectly quantified caspase-1 activation by measuring ASC speck formation. Indeed, single-cell microscopy confirmed that 80 % of cells with GBP1^+^cASp4^+^ STm (indicating active caspase-4), also had ASC specks (active caspase-1) (**Figure 3A**). Notably, caspase-4 was not recruited to ASC specks, which is consistent with previous work (Thurston et al, 2016) (**Figure 3A+B**). As these results suggested dual activation of caspase-1 and caspase-4 in the same cell, we investigated whether and how GBP1 proteolysis might affect caspase-4 recruitment to STm.

**Figure 3:**
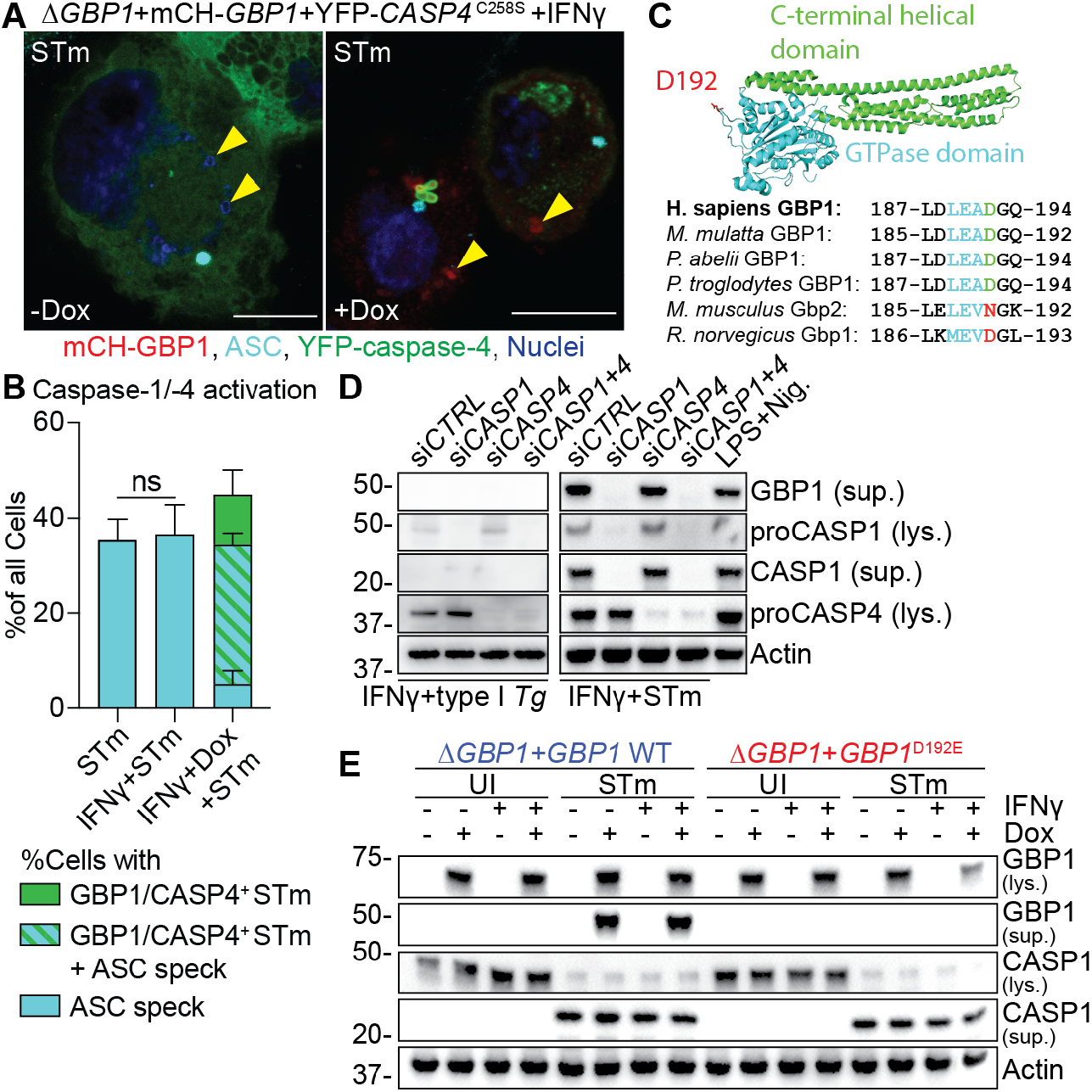
Caspase-1, but not caspase-4, cleaves GBP1 at Asp192 during *Salmonella* infection. **(A)** Representative immunofluorescence images and **(B)** quantification of ASC speck formation and GBP1+caspase-4 recruitment to *Salmonella* Typhimurium (STm) from IFNγ-primed THP-1 Δ*GBP1*+Tet-mCH-*GBP1*+YFP-*CASP4*^C258S^ infected with STm SL1344 (MOI = 30) for 2 hours. Cells were treated with Doxycycline (Dox) to induce GBP1 expression or left untreated. Yellow arrowheads indicate position of some STm within cells (DNA-staining dye). Red: mCH-GBP1; Cyan: ASC; Green: YFP-caspase-4; Blue: Nuclei. Scale bar 10 μm. **(C)** Crystal structure of human GBP1 (PDB: 1F5N) with GTPase domain highlighted in cyan, C-terminal helical domain in green and surface-exposed aspartate D192 in red (top). Multiple sequence alignment of human, primate and rodent GBP1 orthologs depicting the caspase-1 cleavage site (cyan) surrounding Asp192 of human GBP1. **(D)** Representative immunoblots from lysates (lys.) or culture supernatants (sup.) of THP-1 WT, transfected with the indicated siRNA and infected with type I *Toxoplasma gondii* (*Tg*) for 6 hours, STm SL1344 for 4 hours or treated with LPS and Nigericin for 90 minutes. **(E)** Representative immunoblots from lysates or culture supernatants of THP-1 Δ*GBP1*+Tet-*GBP1* WT or *GBP1*^D192E^ cells treated with IFNγ and Dox as indicated and infected with STm SL1344 for 4 hours or left uninfected (UI). **Data information:** Graph in **(B)** shows mean ± SEM of n = 3 independent experiments. *P* values in in **(B)** from two-way ANOVA following adjustment for multiple comparisons; ns, not significant.

We therefore examined the impact of caspase-1-mediated cleavage of GBP1 at the surface exposed Asp192 residue that generates a stable p47 GBP1 C-terminal fragment (**Figure 3C**). Of note, phylogenetic analysis of representative GBPs (Shenoy et al, 2012), revealed that the Asp residue required for caspase-1 cleavage-site was present in all primates and absent in most rodents, including mice (**Figure 3C**). Infection of THP-1 with STm indeed confirmed that GBP1 is cleaved into a ~47 kDa fragment that is detected in supernatants. GBP1 proteolysis could be prevented by silencing caspase-1, but not caspase-4, confirming the dominant role of caspase-1 in the process (**Figure 3D**); LPS+Nigericin treatment served as a positive control and also led to p47 GBP1 production. As expected with the lack of caspase-1 activation during *Tg* infection (Fisch et al, 2019a), GBP1 proteolysis could not be detected in *Tg*-infected THP-1 cells (**Figure 3D**).

To confirm proteolysis of GBP1 at the Asp192 residue, we used a non-cleavable (D192E) variant. We created THP-1 Δ*GBP1* cells expressing the non-cleavable GBP1^D192E^ mutant without or with an mCherry tag (THP-1 Δ*GBP1*+Tet-*GBP1*^D192E^ and THP-1 Δ*GBP1*+Tet-mCH-*GBP1*^D192E^ cells; **Figure S7A**). Immunoblotting of GBP1 from STm-infected IFNγ-primed macrophages revealed caspase-1 activation and formation of p47 GBP1 from cells expressing wildtype GBP1 but not GBP1^D192E^ (**Figure 3E**). Together, these results point towards the specificity of caspase-1 in cleaving GBP1 and that neither caspase-4 nor caspase-8 (active during *Tg* infection) can replace its role.

### Caspase-1-cleaved GBP1 fragments cannot traffic to microbial vacuoles or mediate cell death

As GBP1 can be cleaved by caspase-1, we wanted to investigate how this affects the pathogen-proximal activities of GBP1 in enabling PAMP access and triggering cell death. We infected mCH-GBP1^D192E^ expressing cells with STm and quantified GBP1 recruitment to bacteria. The proportion of GBP1^+^ STm was similar in cells expressing GBP1 WT and GBP1^D192E^ (**Figure 4A**). However, the mean fluorescence intensity of mCH-GBP1 around decorated STm was markedly higher in cells expressing the GBP1^D192E^ variant (**Figure 4A**), even though the expression and fluorescence of GBP1 WT and GBP1^D192E^ was comparable in uninfected cells (**Figure S7B**). In agreement with increased GBP1 amounts covering cytosolic bacteria, STm-infected GBP1^D192E^ cells underwent higher pyroptosis than wildtype cells but released similar levels of IL-1β (**Figure 4B**). This finding is consistent with a major role for GBP1 in promoting caspase-4-driven pyroptosis, but not canonical caspase-1 activation, which is responsible for IL-1β production (Kortmann et al, 2015; Reyes Ruiz et al, 2017). These results led us to speculate that cleavage of GBP1 reduces the cellular pool of functional full-length GBP1, and its cleaved fragments do not support cell death-related roles. Indeed, Δ*GBP1* cells reconstituted with GBP1^1-192^ or GBP1^193-592^ with or without mCherry-tag using our Dox-inducible system (**Figure S7C**) revealed that neither fragment was recruited to STm (**Figure 4C**) nor supported enhanced pyroptosis (**Figure 4D**). As caspase-1 is not activated during *Tg* infection, we anticipated that *Tg* targeting and apoptosis would be similar in cells expressing GBP1 WT or GBP1^D192E^. Indeed, the proportion of *Tg*-PVs decorated with GBP1 WT and GBP1^D192E^ was similar (**Figure 4E**) and apoptosis remained unaffected (**Figure 4F**).

**Figure 4:**
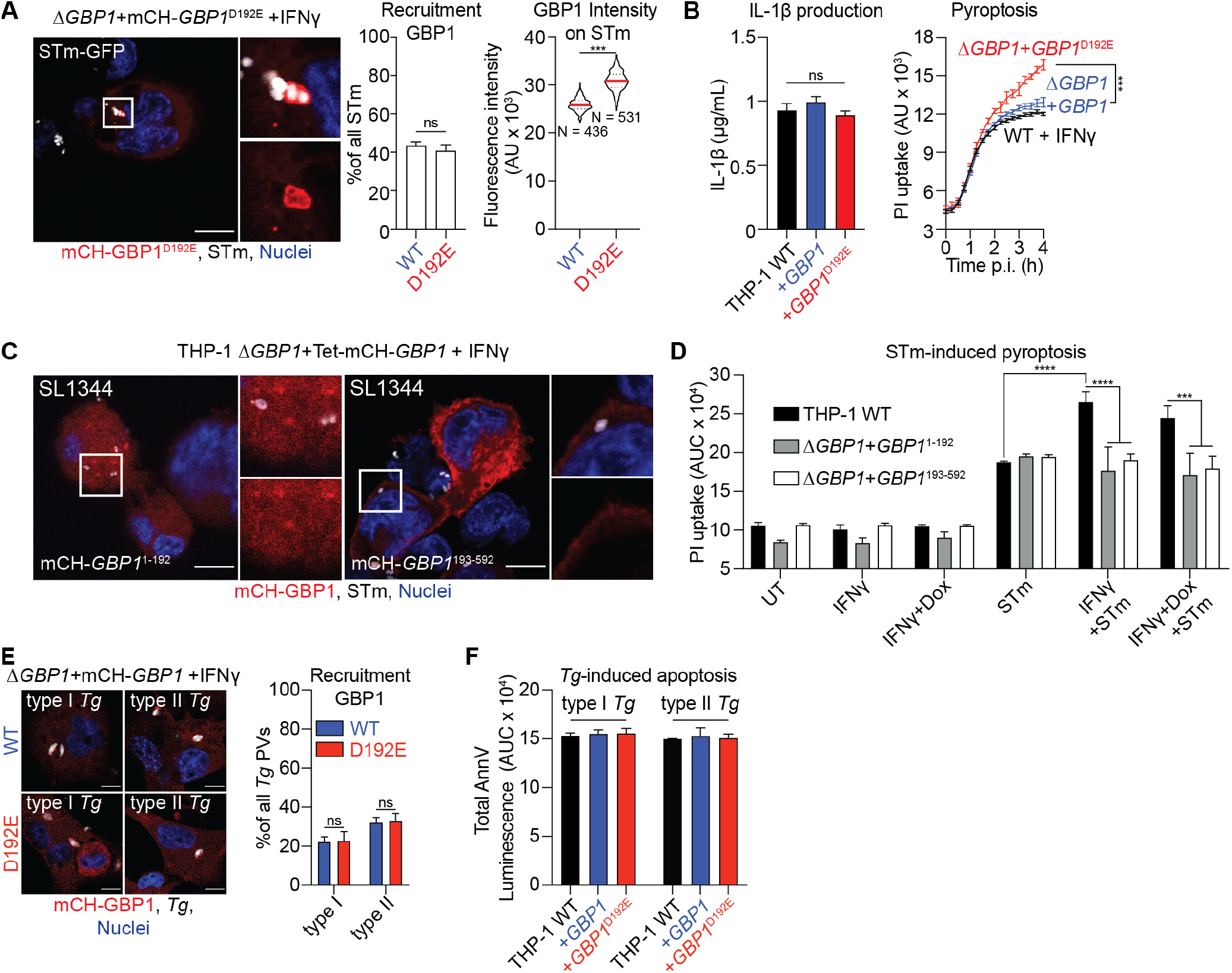
Caspase-1-driven GBP1 proteolysis regulates cell death during *Salmonella*, but not *Toxoplasma*, infection. **(A)** Representative immunofluorescence images and quantification of recruitment and fluorescence intensities of GBP1 to *Salmonella* Typhimurium (STm) in IFNγ-primed and Doxycycline (Dox)-treated THP-1 Δ*GBP1*+Tet-mCH-*GBP1* or mCH-*GBP1*^D192E^ cells infected with STm SL1344-GFP (MOI = 30) for 2 hours. Red: mCH-GBP1; Grey: STm; Blue: Nuclei. Scale bar 10 μm. N depicts the number of measured GBP1+ bacteria in the respective condition. **(B)** Left: IL-1β ELISA from the indicated THP-1 cells primed with IFNγ and Dox and infected with STm SL1344 (MOI = 30) at 4 h post-infection. Right: Real-time propidium iodide (PI) uptake assay from IFNγ-primed THP-1 cells of the indicated genotypes and infected with STm SL1344 for 4 h. **(C)** Representative immunofluorescence image of mCherry tagged GBP1 fragments in IFNγ- and Dox-primed THP-1 *ΔGBP1+Tet* cells expressing the indicated GBP1 fragment infected with STm SL1344 (MOI = 30) for 2 hours. Red: mCH-GBP1; White: STm; Blue: Nuclei. Scale bar 10 μm. **(D)** Area under the curve (AUC) from 4 hours live propidium iodide (PI) uptake cell death assay in THP-1 WT or THP-1 *ΔGBP1+Tet-GBP1* cells expressing either GBP1 fragment 1-192 or 193-592, pre-stimulated with IFNγ only, with IFNγ and Dox to induce GBP1 expression or left untreated (UT) and infected with STm SL1344 (MOI = 30). **(E)** Representative immunofluorescence images and quantification of recruitment of GBP1 in IFNγ-primed and Dox-treated THP-1 Δ*GBP1*+Tet-mCH-*GBP1* or mCH-*GBP1*^D192E^ cells infected with type I or type II *Toxoplasma gondii* (*Tg*) for 6 hours. Red: mCH-GBP1; Grey: Tg; Blue: Nuclei. Scale bar 10 μm. **(F)** AnnV-Glo assay of THP-1 WT and Δ*GBP1* cells stably reconstituted with *Tet-GBP1* WT or *GBP1*^D192E^ as indicated, infected with type I or type II *Tg* for 18 h. Plotted as area under the curve (AUC) from real-time assays. **Data information:** Graphs in **(A)**, **(B)**, **(D)**, **(E)** and **(F)** show mean ± SEM from n = 3 independent experiments. *** *P* ≤ 0.001, **** *P* ≤ 0.0001 for indicated comparisons in **(A)** from nested one-way ANOVA comparing means of the n = 3 experiments (left) or from one-way ANOVA (right), in **(B)** at 4 hours p.i. only and **(D)** from two-way ANOVA and in **(E)** from one-way ANOVA following adjustment for multiple comparisons; ns, not significant.

In summary, these results suggested that active caspase-1 cleaves a portion of cellular GBP1 and generates protein fragments that cannot (1) target cytosolic STm, (2) subsequently recruit caspase-4 and (3) enhance pyroptosis induction. Because IL-1β maturation was not affected by GBP1^D192E^ mutation, we speculate that this caspase-1-driven feedback mechanism balances cell death and IL-1β secretion during STm infection. Moreover, as caspase-1 does not contribute to cell death during *Tg*-infection, this feedback regulatory mechanism is pathogen-specific.

## Discussion

IFNγ-inducible GBPs have emerged as important proteins in host defense against a range of pathogens in the murine system (Meunier & Broz, 2016; Pilla-Moffett et al, 2016; Man et al, 2017; Saeij & Frickel, 2017; Tretina et al, 2019). In this study we have established that human GBP1 is essential for the breakdown of PVMs and *Tg* parasites through the use of two new single-cell assays combined with the artificial intelligence-driven image analysis pipeline HRMAn that are adaptable for other pathogens (Fisch et al, 2019b). In contrast to *Tg*, GBP1 only decorates cytosolic STm and forms a complex with caspase-4, which it recruits onto the surface of bacteria. Caspase-1, but not caspase-4, also cleaves GBP1 at Asp192 to limit pyroptosis. These findings provide important new insights on this key GTPase in human macrophages.

As the forerunner of the human GBP family, GBP1 has been extensively studied structurally and biochemically. For instance, high-resolution structural and biophysical studies point towards an exceptionally fast GTP hydrolysis capacity to form GDP and GMP (Cheng et al, 1991; Schwemmlel & Staeheli, 1994; Praefcke et al, 1999; Ghosh et al, 2006), ability to homo- and hetero-oligomerize (Wehner et al, 2012; Ince et al, 2017; Barz et al, 2019; Lorenz et al, 2019), and undergo isoprenylation (Nantais et al, 1996;

Olszewski et al, 2006; Tripal et al, 2007; Britzen-Laurent et al, 2010). Further, GBP1 is a dynamin-like GTPase and may actively alter biological membranes (Huang et al, 2019); indeed, recombinant farnesylated GBP1 can bend giant unilamellar vesicle membranes in vitro (Shydlovskyi et al, 2017). We add the role of recruiting caspase-4 to STm dependent on functional GTPase activity and isoprenylation, which is in line with previous findings on mouse and human GBPs *in vitro* and *in cellulo* (Nantais et al, 1996; Stickney & Buss, 2000; Prakash et al, 2000; Britzen-Laurent et al, 2010; Fres et al, 2010; Piro et al, 2017; Shydlovskyi et al, 2017; Kohler et al, 2019).

Targeting of *Tg* vacuoles by murine GBPs and their interplay with the IRG proteins has been extensively studied (MacMicking *et al*, 2003; Bernstein-Hanley *et al*, 2006; Singh *et al*, 2006; Degrandi *et al*, 2007; Henry *et al*, 2007; Miyairi *et al*, 2007; Shenoy *et al*, 2007; Coers *et al*, 2008; Hunn *et al*, 2008; Al-Zeer *et al*, 2009; Tiwari *et al*, 2009; Khaminets *et al*, 2010; Lapaquette *et al*, 2010; Singh *et al*, 2010; Brest *et al*, 2011; Traver *et al*, 2011; Virreira Winter *et al*, 2011; Kim *et al*, 2012a; Haldar *et al*, 2013, 2014). Uniquely in mice, two chromosomal loci each encode members of the GBP (~11 genes on Chr 3 and Chr 5) and IRG (~23 genes on Chr 11 and Chr 18) families. Deletion of all mGbps on Chr3 (Δ*Gbp*^Chr3^) abrogates *Tg* vacuole rupture in macrophages and these mice are highly susceptible to *Tg* infection (Yamamoto et al, 2012). Single deletion of mGbp1 (Selleck et al, 2013) or mGbp2 (Degrandi et al, 2013) also results in enhanced susceptibility to *Tg in vivo* and *in vitro.* mGbp2 can homodimerize or form heterodimers with mGbp1 or mGbp5 before recruitment and attack of *Tg* vacuoles (Kravets et al, 2016). However, in the mouse, the hierarchical recruitment of IRG family GTPases to *Tg* vacuoles precedes the recruitment of GBP family members. No GBPs are recruited to *Tg* in *Irgm1/Irgm3*^-/-^ murine cells pointing to their pivotal role in this process (Haldar et al, 2013) In addition to the absence of IRGs in humans, a direct role for individual GBPs in *Tg* vacuole disruption has not been demonstrated for GBPs before, even though mouse Gbp2 has been found to localize inside *Tg* (Kravets et al, 2016). Indeed, while targeting of murine Gbps to *Tg* vacuoles with subsequent vacuolar lysis has been demonstrated for mGbp1, 2 and the collective Gbps located on chromosome 3, parasite plasma membrane lysis has not been observed before (Yamamoto et al, 2012; Degrandi et al, 2013; Selleck et al, 2013).

During STm infection GBP1 only targeted cytosolic bacteria even though a proportion of bacteria remained vacuolar. Our finding that GBP1 only targets bacteria already in the cytosol are consistent with bacterial staining with Gal-8, which binds to glycans on endogenous damaged membranes and recruits other proteins, including the autophagy machinery (Thurston et al, 2012). Furthermore, mouse Gbp-recruitment is reduced in murine macrophages lacking Gal-3, which normally labels *Legionella* (Creasey & Isberg, 2012; Pilla et al, 2014; Feeley et al, 2017; Liu et al, 2018) or *Yersinia* (Feeley et al, 2017) expressing secretion systems that trigger damage of bacterial-containing vacuoles. Work with bacterial mutants that readily access the cytosol, such as *Legionella pneumophila ΔsdhA* and STm *ΔsifA*, also revealed no differences in cytosolic bacteria in mouse Δ*Gbp*^Chr3^ macrophages (Pilla et al, 2014). Similarly, release of *Francisella novicida* into the cytosol was shown to be independent of mouse Gbps (Man et al, 2015; Meunier et al, 2015). It is plausible that in human macrophages GBP1 is dispensable for release of STm into the cytosol even though Gbps encoded at mouse Chr3 and mGbp2 have previously been implicated in this process in murine cells (Meunier et al, 2014). It is tempting to speculate that human GBP1 recruitment to vacuolar STm is prevented by a bacterial virulence factor. Indeed, anti-GBP1 bacterial effectors have been identified in *Shigella flexneri* (Li et al, 2017; Piro et al, 2017; Wandel et al, 2017). Further work should investigate whether other human GBPs also assemble alongside or assist GBP1 during STm infection.

Our work also shows unique GBP1 action during infection by these diverse pathogens whose distinct PAMPs are recognized by downstream innate immune pathways. Click-chemistry revealed that *Tg*-DNA is present in the cytoplasm of GBP1-expressing macrophages that subsequently induces the assembly of the atypical AIM2-ASC-caspase-8 SMOC and apoptosis. Super-resolution imaging structure of a caspase-8 containing AIM2 inflammasome closely resembles previously published structures of caspase-8 in NLRP3/NLRC4 inflammasomes (Man et al, 2013, 2014), revealing donut-like ASC rings enclosing AIM2 and caspase-8. Super-resolution microscopy during STm infection showed that GBP1 and caspase-4 formed a dense coat on STm, which reduced bacterial staining with anti-LPS antibody. Whether this was due to reduced antibody access due to the GBP1/caspase-4 coat or blocking of the LPS epitope by caspase-4 cannot be definitively distinguished. As caspase-4 by itself could not recruit to the bacteria, we speculate that GBP1 is involved in exposing parts of the LPS that are buried deeper within the membrane potentially through direct interaction with LPS as has been suggested for mGbp5 (Santos et al, 2018). We therefore hypothesize that GBP1 ‘opens’ the bacterial outer membrane for caspase-4 to gain access to otherwise hidden PAMPs.

Our results also uncovered a physiological role for GBP1 proteolysis by caspase-1 that was previous reported in vitro using HUVEC cells and *in vivo* from cerebrospinal fluid of meningitis patients (Naschberger et al, 2017), which we confirmed during natural infection of macrophages with STm. Not surprisingly, GBP1 is not proteolyzed during *Tg* infection due to the absence of active caspase-1 in this setting (Fisch et al, 2019a). Notably, despite the 40-98 % sequence similarity between human and mouse GBPs (Shenoy et al, 2007; Kim et al, 2011), and the conservation of Asp192 in other primate GBP1 sequences, Asp192 is absent in the closest murine homologue, mGbp2 (Olszewski et al, 2006), which is therefore unlikely to be regulated in this manner. Intriguingly, this finding mirrors our recent identification of the proteolysis of human, but not mouse p62, by caspase-8 at a conserved residue found in other mammalian p62 sequences (Sanchez-Garrido et al, 2018). During STm infection, caspase-1 plays a dominant role in IL-1β maturation whereas IFNγ-induced GBP1 enhances caspase-4-driven pyroptosis. As a result, caspase-1-dependent proteolysis of GBP1 affected pyroptosis but not IL-1β maturation. Furthermore, GBP1 fragments produced by caspase-1 failed to target STm, recruit caspase-4 and support pyroptosis. Thus, besides directly aiding the release or access to PAMPs for detection by caspases, GBP1 itself is a target of caspase-1 and a key regulatory hub that modulates host cell death. This contrasts our discovery of the ubiquitin conjugating enzyme UBE2L3 as an indirect target of caspase-1 that specifically controls IL-1β production but not pyroptosis (Eldridge et al, 2017). At the whole organism level, these mechanisms potentially enable differential responses based on the strength of the activating stimulus. Enhanced IL-1β or IL-18 production for adaptive immunity may be balanced by cell death that could enable pathogen uptake by other cell types such as neutrophils. Studies on cellular targets of caspases may therefore provide new insights on homeostasis and disease.

Common themes also emerge from work on human and mouse GBPs. For instance, human GBP1 and mouse Gbp2 accumulate on vesicles generated through sterile damage, which suggests they could detect endogenous luminal ligands in the cytosol, for example endogenous sulfated lipids (Bradfield, 2016). The presence of Gal-3/Gal-8 and GBPs at sites of damaged membranes suggests GBPs may be assisted in sensing damage by other proteins, including IFN-induced genes. Undoubtedly, future work in the area will focus on finding how human GBPs are targeted to diverse microbes, the ligands they sense and how they are regulated.

## Author Contributions

DF, ARS and EMF conceived the idea for the study, DF, BC, MCD and VE performed experiments. HB, MY provided essential reagents. LMC and APS provided essential expertise and equipment. DF, ARS and EMF analyzed and interpreted the data and wrote the manuscript. All authors revised the manuscript.

## Acknowledgements

We would like to thank Matt Renshaw from the Crick Advanced Light Microscopy (CALM) STP for help with super-resolution SIM imaging, Julia Sanchez-Garrido for help in optimizing immunoblots and advice on reagents, Michael Howell from the Crick High-throughput screening (HTS) STP for help in performing automated imaging experiments, the Crick Genomics and Equipment Park STP for performing Sanger sequencing and DNA minipreps for cloning, Debipriya Das from the Crick Flow Cytometry STP for sorting Tg parasites and Caia Dominicus, Jeanette Wagener and Joanna Young from Moritz Treeck’s lab for help with creation of new transgenic Tg lines. We thank all members of the Frickel and the Shenoy labs for productive discussion and Crick Core facilities for assistance in the project.

## Materials And Methods

### Cells, parasites and treatments

THP-1 (TIB-202, ATCC) were maintained in RPMI with GlutaMAX (Gibco) supplemented with 10% heat-inactivated FBS (Sigma), at 37°C in 5% CO2 atmosphere. THP-1 cells were differentiated with 50 ng/mL phorbol 12-myristate 13-acetate (PMA, P1585, Sigma) for 3 days followed by a rested for 2 days in complete medium without PMA. Cells were not used beyond passage 20. HEK293T and human foreskin fibroblasts (HFF) were maintained in DMEM with GlutaMAX (Gibco) supplemented with 10% FBS at 37°C in 5% CO2 atmosphere. *Tg* expressing luciferase/eGFP (RH type I and Prugniaud (Pru) type II) were maintained by serial passage on monolayers of HFF cells. All cell culture was performed without addition of antibiotics unless otherwise indicated. Cell lines were routinely tested for mycoplasma contamination by PCR and agar test. An overview of all cell lines made/ used in this study is provided in **Table S1**.

Cells were stimulated for 16 h prior to infection in complete medium at 37°C with addition of 50 IU/mL human IFNγ (285-IF, R&D Systems). Induction of Doxycycline-inducible cells was performed with 200 ng/mL Doxycycline overnight (D9891, Sigma). To chemically activate caspase-1, cells were treated with 10 μM Nigericin (N1495, Invitrogen) and 100 μg/mL LPS-Sm (IAX-100-011, Adipogen).

### *Toxoplasma gondii* infection

Parasite were passaged the day before infection. *Tg* tachyzoites were harvested from HFFs by scraping and syringe lysing the cells through a 25 G needle. The *Tg* suspension was cleared by centrifugation at 50 x g for 5 min and then the parasites were pelleted by centrifugation at 550 x g for 7 min from the supernatant, washed with complete medium, and finally resuspended in fresh medium. Viable parasites were counted with trypan blue and used for infection at a multiplicity of infection (MOI) of 3 for most experiments or 1 for immunofluorescence imaging. Infection was synchronized by centrifugation at 500 x g for 5 min. Two hours after infection, extracellular parasites were removed with three PBS washes.

### Flow cytometry and sorting

For flow cytometry analysis of GFP-fluorescence, *Tg* Δ*Hpt*+GFP_1-10_ were harvested from host cells by syringe lysis, washed twice with warm PBS and then re-suspended in PBS + 1% BSA. Parasites were analyzed on a LSR Fortessa (BD Biosciences) and data were processed using FlowJo version 10.3 (FlowJo, LLC). For viability determination of GFP-fluorescing versus non-fluorescing *Tg* the parasites were harvested and prepared identically, sorted on a FACSAria™ III (BD Biosciences) based on their GFP-fluorescence and then plated onto HFFs grown confluent in wells of a 24-well plate. 5 days post infection of the HFFs, cells were fixed with ice-cold methanol and stained with crystal violet. Following 5 washes with PBS, plaques were imaged on a GelCount™ Colony Counter (Oxford Optronix) and cell covered area determined using FIJI ImageJ.

### *Salmonella* Typhimurium infection

STm SL1344-GFP (with pFPV25.1 plasmid) was maintained under Ampicillin selection (11593027, Gibco). STm SL1344 wildtype strain was maintained in the presence of streptomycin (11860038, Gibco) selection. One day before infection bacteria from a single colony were inoculated into 9 mL LB and grown overnight at 37°C. The overnight culture was diluted 1:50 into LB + 300 mM NaCl (746398, Sigma) and grown shaking in a closed container until an OD_600_ of 0.9. Bacteria were harvested by centrifugation at 1000 x g for 5 min, washed with serum-free cell culture medium twice and re-suspended in 1 mL medium. Cells were infected with STm at an MOI of 30 and infections were synchronized by centrifugation at 750 x g for 10 min. Infected cells were washed 30 min post-infection three times with warm PBS (806552, Sigma) to remove extracellular bacteria and fresh medium containing 100 μg/mL Gentamicin (15750060, Gibco) was added for 1 h. Medium was then replaced with medium containing 10 μg/mL gentamicin and the infection continued for indicated times. Bacterial MOI used for infections were confirmed by plating on LB agar plates.

### Creation of transgenic *Toxoplasma gondii*

To create new *Tg* lines that constitutively express non-fluorescent GFP_1-10_ fragment, the GFP_1-10_ ORF was amplified from pEGFP-C1 (Clontech) and Gibson-assembled into NsiI and PacI digested pGRA-HA-HPT (a gift from Moritz Treeck) (Coppens et al, 2006), to have expression of the ORF under control of the *Tg*GRA1 promoter.

Next the plasmid was transfected into type II (Pru) *Tg* Δ*Hpt* (a gift from Moritz Treeck) using nucleofection as established by Young et al (Young et al, 2019): The plasmid was linearized using PsiI-V2 and purified using phenol-chloroform precipitation and resuspended in P3 solution (Lonza). Successful linearization was confirmed using agarose-gel electrophoresis. Next, type II (Pru) *Tg* Δ*Hpt* were harvested from HFFs by syringe lysis and washed with PBS twice and then 5×10^6^ parasites resuspended in P3 solution. Prior to nucleofection, 25 μg linearized DNA were added to the parasites and then nucleofected using 4D-NucleofectorTM (Lonza) with setting EO-115. Transfected parasites were then incubated for 12 minutes at room temperature, followed by platting onto fresh HFF cells into a T25 tissue culture flask. The next day, medium was replaced with complete DMEM containing 50 μg/mL xanthine and mycophenolic acid (MPA) each for selection. The selection medium was replaced every two days and the parasites passaged normally for two weeks when all *Tg* in the untransfected control had died. Successful integration of the plasmid and expression of GFP_1-10_ was confirmed by immunofluorescence and immunoblotting.

### Creation of new cell lines

#### Creation of the Dox-inducible GBP1 and caspase-8 cell lines

THP-1 Δ*GBP1*+Tet-*GBP1* and THP-1 Δ*GBP1*+Tet-mCH-*GBP1* have been published before and the THP-1 WT+Tet-*CASP8*-Flag cells were created identically using Lentiviral transductions (Fisch et al, 2019a).

To create the caspase-8-3xFlag expressing Dox-inducible plasmid (pLenti-Tet-*CASP8*-3xFlag), the empty vector backbone was digested with BamHI, *CASP8* ORF was amplified from pcDNA3-*CASP8* by PCR (Addgene #11817, a gift from Guy Salvesen) (Stennicke & Salvesen, 1997), 3xFlag was amplified from lentiCRISPRv2 (Addgene #52961, a gift from Feng Zhang) (Sanjana et al, 2014) and all fragments assembled with a Gibson assembly. Similarly, to create the 3xFlag-*GBP*1 expressing Dox-inducible plasmid (pLenti-Tet-3xFlag-*GBP1*), the empty vector backbone was digested with BamHI, *GBP1* ORF was amplified from pGene-*GBP1* by PCR (Frickel lab), 3xFlag was amplified from lentiCRISPRv2 and all fragments assembled with a Gibson assembly. In the same way, GBP1 fragments 1-192 and 193-592 were amplified from pGene-*GBP1* (Frickel lab) and Gibson assembled into BamHI digested pLenti-Tet (Fisch et al, 2019a) with and without addition of a mCherry tag. The obtained plasmids were then transduced into THP-1 Δ*GBP1*+Tet cells (Fisch et al, 2019a) using lentiviral particles.

To make the cells expressing GBP1^D192E^, the pLenti-Tet-mCH-*GBP1* and pLenti-Tet-*GBP1* plasmids (Fisch et al, 2019a) were mutated using site-directed mutagenesis and transduced into the THP-1 Δ*GBP1*+Tet target cells using Lentiviral transduction as described before (Fisch et al, 2019a). To make cells expressing YFP-*CASP4*^C258S^ and mutated GBP1 versions, THP-1 Δ*GBP1*+Tet-mCH-*GBP1*^K51A^, +Tet-mCH-*GBP1*^C589A^ or +Tet-mCH-*GBP1*^Δ589-592^ (Fisch et al, 2019a) were transduced with pMX-CMV-YFP-*CASP4*^C258S^ (Fisch et al, 2019a) using retroviral particles. All primers used for cloning PCRs can be found in **Table S3**.

#### Creation of myc-AIM2 expressing cell line

To create a lentiviral vector for constitutive expression of myc-AIM2, the ORF was amplified from pcDNA3-*myc-AIM2* (Addgene #73958, a gift from Christian Stehlik) (Khare et al, 2014) and Gibson assembled into BstBI and BsrGI digested pLEX-MCS-*ASC*-GFP (Addgene #73957, a gift from Christian Stehlik) (de Almeida et al, 2015) to create pLEX-MCS-myc-*AlM2*. The newly made vector was then transduced into THP-1 +Tet-*CASP8*-Flag cells to create THP-1+Tet-*CASP8* + myc-*AIM2* cells using Lentiviral transduction as described above.

#### Creation of GFP_11_ expressing cell lines

To create an lentiviral vector for constitutive expression of GFP_11_, the sgRNA cassette from lentiCRISPRv2 was removed by digestion with KpnI and EcoRI and the plasmid re-ligated using annealed repair oligo pair 1 (see **Table S3**) and Quick Ligation™ Kit (M2200L, NEB). Next, the Cas9-ORF was removed by digestion with XbaI and BamHI and again the vector re-ligated using annealed repair oligo pair 2 (see **Table S3**), also adding a multiple cloning site, which created the vector pLenti-P2A-Puro. Next, the GFP_11_ ORF was amplified from pEGFP-C1 (Clontech) and ligated into BamHI and XbaI digested pLenti-P2A-Puro, to have the GFP_11_-ORF in frame with the P2A-Puro cassette, for Puromycin-selectable, constitutive expression of GFP_11_. The newly made vector was then transduced into THP-1 WT and THP-1 Δ*GBP1*+Tet-*GBP1* cells using Lentiviral transduction as described above.

#### Real-time cell death assays and IL-1β ELISA

To measure live kinetics of cell death, 60,000 cells were seeded per well of a black-wall, clear-bottom 96-well plate (Corning) for differentiation with PMA, treated and infected as described above. Medium was replaced with phenol-red-free RPMI supplemented with 5 μg/mL propidium iodide (P3566, Invitrogen). The plate was sealed with a clear, adhesive optical plate seal (Applied Biosystems) and placed in a plate reader (Fluostar Omega, BMG Labtech) pre-heated to 37°C. PI fluorescence was recorded with top optics every 15 min for times as indicated.

Apoptosis kinetics were analyzed using the RealTime-Glo™ Annexin V Apoptosis Assay (JA1001, Promega) according to the manufacturer’s instruction. Simultaneously with infection, detection reagent was added. Luminescence was measured using a Fluostar Omega plate reader (BMG Labtech). No-cell, medium-only controls were used for background correction.

For IL-1β ELISA, the cell culture supernatant was harvested, cleared by centrifugation at 2000 x g for 5 minutes and diluted in the buffer provided with the ELISA kit. ELISA was performed according to the manufacturer’s instruction. IL-1β ELISA kit was from Invitrogen (#88-7261, detection range 2 - 150 pg/mL).

### Immunoblotting and gel staining

For immunoblotting, 0.5×10^6^ cells were seeded per well of a 48-well plate, differentiated with PMA, pre-treated and infected. Cells were washed with ice-cold PBS and lysed for 5 min on ice in 50 μL RIPA buffer supplemented with protease inhibitors (Protease Inhibitor Cocktail set III, EDTA free, Merck) and phosphatase inhibitors (PhosSTOP, Roche). Lysates were cleared by centrifugation at full speed for 15 min at 4°C. BCA assay (Pierce BCA protein assay kit, 23225, Thermo Scientific) was performed to determine protein concentration. 10 μg of total protein per sample were run on Bis-Tris gels (Novex, Invitrogen) in MOPS running buffer and transferred on Nitrocellulose membranes using iBlot transfer system (Invitrogen). Membranes were blocked with either 5% BSA (A2058, Sigma) or 5% dry-milk (M7409, Sigma) in TBS-T for at least 1 h at room temperature. Incubation with primary antibodies was performed at 4°C overnight (all antibodies used in this study can be found in **Table S2**). Blots were developed by washing the membranes with TBS-T, probed with 1:5000 diluted secondary antibodies in 5% BSA in TBS-T and washed again. Finally, the membranes were incubated for 2 minutes with ECL (Immobilon Western, WBKLS0500, Millipore) and luminescence was recorded on a ChemiDoc MP imaging system (Biorad). For silver staining of protein gels, following SDS-PAGE, the gels were washed in ddH2O and then silver stained following the manufacturers instruction (Silver Stain Plus Kit, 1610449, Biorad).

For immunoblots of culture supernatants, cells were treated in OptiMEM (1105802, Gibco) without serum. Proteins in the supernatants were precipitated with 4x volume cold acetone (V800023, Sigma) overnight at −20°C, and pelleted by centrifugation. Pellets were air dried and re-suspended in 40 μL 2x Laemmli loading dye.

### Identification of GBP1 interacting proteins by mass spectrometry

#### Sample preparation

10×10^6^ THP-1 Δ*GBP1*+Tet-Flag-*GBP1* cells were seeded in 6-well plates and differentiated, pre-treated with IFNγ and Doxycycline and infected with STm as described before. 2 hours p.i. the interacting proteins were crosslinked with 1% formaldehyde (28906, Thermo Scientific) for 10 minutes at room temperature and the reaction quenched by addition of 125 mM glycine (Sigma). Cell were washed in ice-cold PBS and scraped from the plates. Cells were then pelleted by centrifugation and washed in PBS. Whole-cell lysates were prepared by adding 500 uL lysis buffer (1% Triton X-100, 20 mM Tris-HCl [pH 8], 130 mM NaCl, 1 mM dithiothreitol, 10 mM sodium fluoride, protease inhibitors (Protease Inhibitor Cocktail set III, EDTA free, Merck), phosphatase inhibitor cocktails (PhosSTOP, Roche)) and incubation for 15 minutes on ice. Lysates were cleared by centrifugation and then added to Flag(M2)-agarose beads (A2220, Sigma) washed three times with lysis buffer. Flag-GBP1 was captured by incubation on a rotator overnight at 4°C. Beads were then washed once with lysis buffer, three times with lysis buffer containing 260 mM NaCl and then again twice with lysis buffer. Proteins were eluted using 200 μg/mL 3xFlag peptide (F4799, Sigma) in lysis buffer by incubation on an orbital shaker (1400 rpm) for 2 hours at room temperature. Samples were then prepared by adding loading dye containing 5% β-Mercaptoethanol (Sigma) to reverse crosslinking and run on a 12% Bis-Tris polyacrylamide gel until the running front had entered the gel roughly 5 mm.

#### Trypsin digestion

Samples on the SDS-PAGE were excised as three vertical lanes each. The excised gel pieces were destained with 50% acetonitrile/50 mM ammonium bicarbonate, reduced with 10 mM DTT, and alkylated with 55 mM iodoacetamide. After alkylation, the proteins were digested with 250 ng of trypsin overnight at 37°C. The resulting peptides were extracted in 2% formic acid, 1% acetonitrile and speed vacuum dried. Prior to analysis the peptides were reconstituted in 50 μl of 0.1% TFA.

#### Mass spectrometry

The peptides were loaded on a 50 cm EASY-Spray™ column (75 μm inner diameter, 2 μm particle size, Thermo Fisher Scientific), equipped with an integrated electrospray emitter. Reverse phase chromatography was performed using the RSLC nano U3000 (Thermo Fisher Scientific) with a binary buffer system at a flow rate of 275 nL/min. Solvent A was 0.1% formic acid, 5% DMSO, and solvent B was 80% acetonitrile, 0.1% formic acid, 5% DMSO. The in-gel digested samples were run on a linear gradient of solvent B (2 - 30%) in 95.5 min, total run time of 120 min including column conditioning. The nano LC was coupled to an Orbitrap Fusion Lumos mass spectrometer using an EASY-Spray™ nano source (Thermo Fisher Scientific). The Orbitrap Fusion Lumos was operated in data-dependent acquisition mode acquiring MS1 scan (R=120,000) in the Orbitrap, followed by HCD MS2 scans in the Ion Trap. The number of selected precursor ions for fragmentation was determined by the “Top Speed” acquisition algorithm with a cycle time set at 3 seconds. The dynamic exclusion was set at 30s. For ion accumulation the MS1 target was set to 4×10^5^ ions and the MS2 target to 2×10^3^ ions. The maximum ion injection time utilized for MS1 scans was 50 ms and for MS2 scans was 300 ms. The HCD normalized collision energy was set at 28 and the ability to inject ions for all available parallelizable time was set to “true”.

#### Data processing and analysis

Orbitrap .RAW files were analyzed by MaxQuant (version 1.6.0.13), using Andromeda for peptide search. For identification, peptide length was set to 7 amino acids, match between runs was enabled and settings were kept as default. Parent ion and tandem mass spectra were searched against UniprotKB *Homo sapiens* and *Salmonella* Typhimurium databases. For the search the enzyme specificity was set to trypsin with maximum of two missed cleavages. The precursor mass tolerance was set to 20 ppm for the first search (used for mass re-calibration) and to 6 ppm for the main search. Product mass tolerance was set to 20 ppm. Carbamidomethylation of cysteines was specified as fixed modification, oxidized methionines and N-terminal protein acetylation were searched as variable modifications. The datasets were filtered on posterior error probability to achieve 1% false discovery rate on protein level. Quantification was performed with the LFQ algorithm in MaxQuant using three replicate measurements per experiment.

### Quantitative RT-PCR (qRT-PCR)

RNA was extracted from 0.25×10^6^ cells using Trizol reagent (15596026, Invitrogen). RNA (1 μg) was reverse transcribed using high-capacity cDNA synthesis kit (4368813, Applied Biosystems). qPCR used PowerUP SYBR green (A25742, Applied Biosystems) kit, 20 ng cDNA in a 20 μL reaction and primers (all primer used for qPCR can be seen in **Table S3**) at 1 μM final concentration on a QuantStudio 12K Flex Real-Time PCR System (Applied Biosystems). Recorded Ct values were normalized to Ct of human HPRT1 and data plotted as ΔCt (Relative expression).

### siRNA transfection

Cells were transfected with siRNAs two days prior to infection, at the same time the THP-1 differentiation medium was replaced with medium without PMA. All siRNAs were used at a final concentration of 30 nM. For transfection, a 10x mix was prepared in OptiMEM containing siRNA(s) and TransIT-X2 transfection reagent (MIR 600x, Mirus) in a 1:2 stoichiometry. All siRNAs used in this study can be found in **Table S3**.

### Fixed immunofluorescence microscopy

For confocal imaging 0.25×10^6^ THP-1 cells were seeded on gelatin-coated (G1890, Sigma) coverslips in 24-well plates. Following differentiation, treatments and infection, cells were washed three times with warm PBS prior to fixation to remove any uninvaded pathogens and then fixed with 4% methanol-free formaldehyde (28906, Thermo Scientific) for 15 min at room temperature. Following fixation, cells were washed again with PBS and kept at 4°C overnight to quench any unreacted formaldehyde. Fixed specimens were permeabilized with PermQuench buffer (0.2% (w/v) BSA and 0.02% (w/v) saponin in PBS) for 30 minutes at room temperature and then stained with primary antibodies for one hour at room temperature. After three washes with PBS, cells were incubated with the appropriated secondary antibody and 1 μg/mL Hoechst 33342 (H3570, Invitrogen) diluted in PermQuench buffer for 1 hour at room temperature. Cells were washed with PBS five times and mounted using 5 μL Mowiol.

Specimens were imaged on a Leica SP5-inverted confocal microscope using 100x magnification and analyzed using LAS-AF software. For structured-illumination super-resolution imaging, specimens were imaged on a GE Healthcare Lifesciences DeltaVision OMX SR imaging system and images reconstructed using the DeltaVision software. All images were further formatted using FIJI software. 3D rendering of image stacks and distance measurements of AIM2-ASC-CASP8 inflammasome specks was performed using Imaris 8.3.1.

### Extended microscopy sample preparation and image analysis

#### Quantification of protein recruitment to pathogen vacuoles and ASC speck formation

Specimens were prepared as described above. Images were acquired using a Ti-E Nikon microscope equipped with LED-illumination and an Orca-Flash4 camera using a 60x magnification. All intracellular parasites/bacteria of 100 fields of view were automatically counted based on whether they showed recruitment of the protein of interest using HRMAn high-content image analysis (Fisch et al, 2019b). Further, the analysis pipeline was used to measure the fluorescence intensity of GBP1 on STm vacuoles using the radial intensity measurement implemented in HRMAn.

For quantification of ASC speck formation, 100 *Tg*-infected cells were manually counted per condition using a Ti-E Nikon microscope equipped with LED-illumination using 60x magnification based on whether they contain an ASC speck and whether STm was decorated with GBP1/CASP4. The experiment was repeated independently three times.

#### EdU labeling for visualization of Tg-DNA release

Type I (RH) *Tg* were grown in fully confluent and nonreplicating HFFs for 3 days in the presence of 20 μM EdU to incorporate the nucleotide into their DNA. Labelled parasites were then harvested and used for infection as described above. 6 hours p.i. cells were fixed and EdU incorporated into *Tg*-DNA visualized by staining the specimens using Click-iT™ EdU Cell Proliferation Kit for Imaging, Alexa Fluor™ 647 dye (C10340, Invitrogen) according to the manufacturers instruction. Coverslips were further stained with Hoechst and mounted before imaging on a Ti-E Nikon microscope equipped with LED-illumination and an Orca-Flash4 camera using a 100x magnification.

#### Correlative light and electron microscopy

1.25×10^6^ THP-1 cells were seeded in a μ-Dish35 mm, high Glass Bottom Grid-500 (81168, ibidi) and differentiated with PMA as described before. Cells were then pre-stimulated with IFNγ overnight and infected with type II (Pru) *Tg* at an MOI =1 for 6 hours. One hour prior to fixation, 1 μg/mL CellMask Deep Red (H32721, Invitrogen) and 20 μM Hoechst 33342 (H3570, Invitrogen) was added to the culture medium to label the cells for detection in fluorescence microscopy. Cells were fixed by adding warm 8% (v/v) formaldehyde (Taab Laboratory Equipment Ltd) in 0.2 M phosphate buffer (PB) pH 7.4 directly to the cell culture medium (1:1) for 15min. The samples were then washed and imaged in PB using a Zeiss AiryScan LSM 880 confocal microscope. Samples were then processed using a Pelco BioWave Pro+ microwave (Ted Pella Inc) and following a protocol adapted from the National Centre for Microscopy and Imaging Research protocol (Deerinck et al, 2010) (See **Table S4** for full BioWave program details). Each step was performed in the Biowave, except for the PB and water wash steps, which consisted of two washes followed by two washes in the Biowave without vacuum (at 250 W for 40 s). All chemical incubations were performed in the Biowave for 14 min under vacuum in 2 min cycles alternating with/without 100 W power. The SteadyTemp plate was set to 21°C unless otherwise stated. In brief, the samples were fixed again in 2.5% (v/v) glutaraldehyde (Taab) / 4% (v/v) formaldehyde in 0.1 M PB. The cells were then stained with 2% (v/v) osmium tetroxide (Taab) / 1.5% (v/v) potassium ferricyanide (Sigma), incubated in 1% (w/v) thiocarbohydrazide (Sigma) with SteadyTemp plate set to 40°C, and further stained with 2% osmium tetroxide in ddH_2_O (w/v). The cells were then incubated in 1% aqueous uranyl acetate (Agar Scientific), and then washed in dH2O with SteadyTemp set to 40°C for both steps. Samples were then stained with Walton’s lead aspartate with SteadyTemp set to 50°C and dehydrated in a graded ethanol series (70%, 90%, and 100%, twice each), at 250 W for 40 s without vacuum. Exchange into Durcupan ACM^®^ resin (Sigma) was performed in 50% resin in ethanol, followed by 4 pure Durcupan steps, at 250 W for 3 min, with vacuum cycling (on/off at 30 s intervals), before embedding at 60°C for 48 h. Blocks were serial sectioned using a UC7 ultramicrotome (Leica Microsystems) and 70 nm sections were picked up on Formvar-coated slot copper grids (Gilder Grids Ltd). Consecutive sections were viewed using a 120 kV Tecnai G2 Spirit transmission electron microscope (Thermo Fischer Scientific) and images were captured using an Orius CCD camera (Gatan Inc). Individual TEM images of ~25-30 consecutive sections per *Tg* parasite were converted as Tiff in Digital Micrograph (Gatan Inc.) and aligned using TrakEM2, a plugin of the FIJI framework (Cardona et al, 2012). The stacks were used to check the integrity of the PV and for coarse alignment with the AiryScan data.

#### Vacuole breakage assay (HRMAn)

For quantification of *Tg* vacuole integrity, cells seeded in black-wall 96-well imaging plates were infected and treated as described before. One hour prior to fixation, 1 μg/mL CellMask Deep Red (H32721, Invitrogen) was added to the culture medium to load the cytosol of host cells with this fluorescent dye. Following fixation and staining with Hoechst (H3570, Invitrogen), plates were imaged at 40x magnification on a Cell Insight CX7 High-Content Screening (HCS) Platform (Thermo Scientific) and 25 fields of view per well were recorded. Fluorescence of the dye within detected *Tg* vacuoles was then analyzed using HRMAn (Fisch et al, 2019b). Additionally, HRMAn was used to classify *Tg* vacuoles based on recruitment of mCH-GBP1 using the implemented neural network and the dataset stratified into decorated and non-decorated vacuoles.

#### Differential stain for detection of cytosolic STm

To distinguish between STm contained in vacuoles and bacteria that had escaped into the cytosol of infected macrophages, cells were differentially permeabilized using 25 μg/mL digitonin for one minute at room temperature as has been described before (Meunier & Broz, 2015). Cytosolic STm were then stained using anti-*Salmonella* antibody (ab35156, Abcam) that has been pre-labelled using Alexa Fluor™ 647 Protein Labeling Kit (A20173, Invitrogen) for 15 minutes at 37°C, prior to immediate fixation with 4% paraformaldehyde. Following fixation cells were permeabilized as described above and all STm were stained using the same but unlabeled antibody and corresponding Alexa Fluor 488 labelled secondary antibody. Cells were further stained with Hoechst (H3570, Invitrogen) and imaged on a Leica SP5-inverted confocal microscope using 100x magnification. For quantification, 100 fields of view per coverslip (typically >1000 individual STm overall) were acquired using a Ti-E Nikon microscope equipped with LED-illumination and an Orca-Flash4 camera at 60x magnification and analyzed with HRMAn (Fisch et al, 2019b) for colocalization of fluorescent signal of all and cytosolic STm.

### Data handling and statistics

Data analysis used nested t-test, one-way ANOVA or two-way ANOVA as groups that were compared are indicated in the figure legends. Benjamini, Krieger and Yekutieli false-discovery rate (Q = 5%) based correction for multiple comparisons as implemented in Prism was used when making multiple comparisons. Graphs were plotted using Prism 8.1.1 (GraphPad Inc.) and presented as means of n = 3 experiments (with usually 3 technical repeats within each experiment) with error bars representing SEM, unless stated otherwise. Structure image of GBP1 was created using MacPymol v.1.74.

## Data availability

All datasets generated during and/or analyzed during the current study are available from the corresponding author on request.

## Supplementary Information

**Fig. S1:**
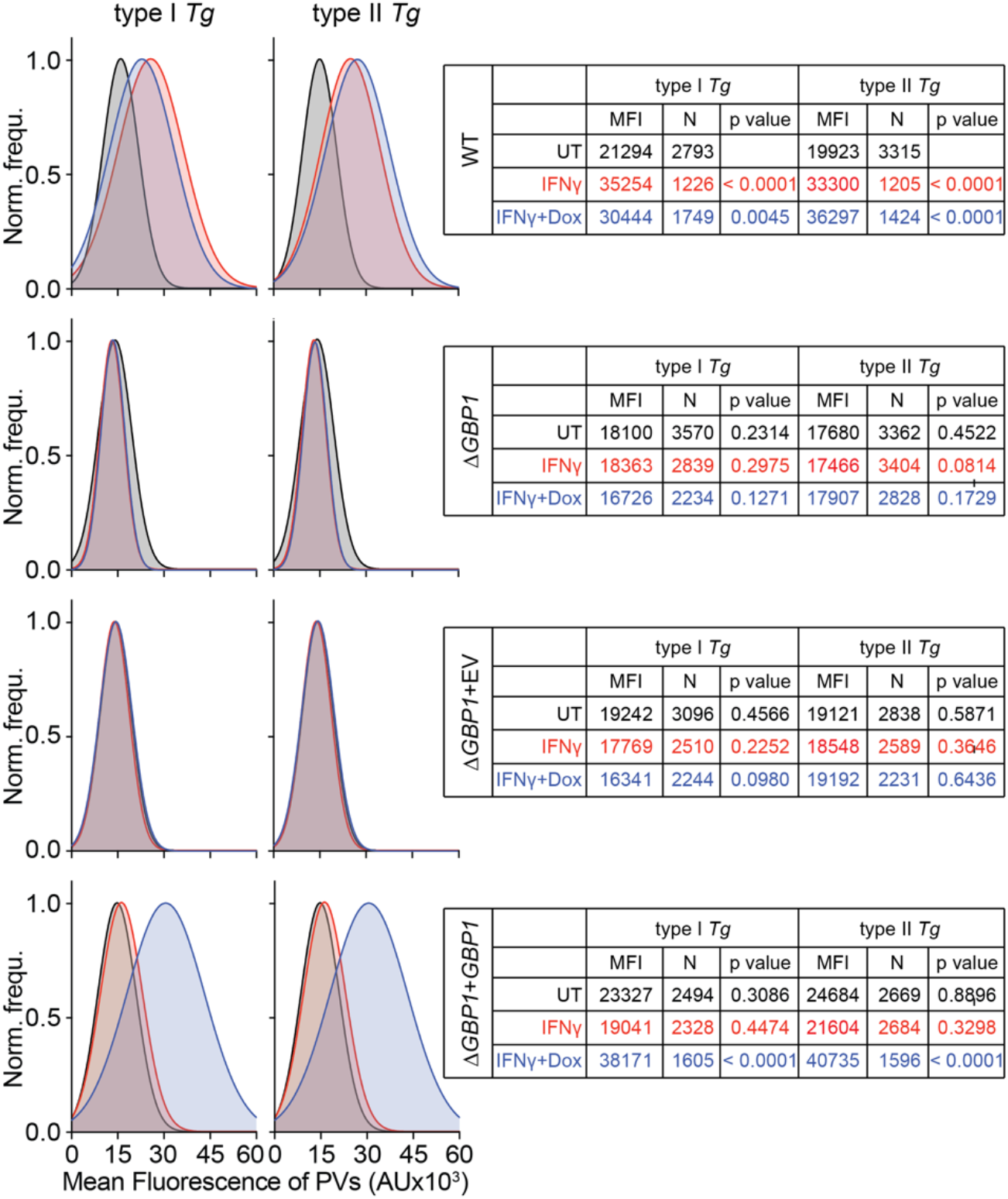
A novel dye influx assay reveals GBP1 contribution to *Toxoplasma* vacuole disruption. Representative normalized frequency plots (Norm. frequ.) and data tables of fluorescence intensities of vacuoles in type I or type II *Toxoplasma gondii* (*Tg*)-infected THP-1 WT, THP-1 *ΔGBP1*, THP-1 Δ*GBP1*+Tet-empty vector (EV) or THP-1 Δ*GBP1*+Tet-*GBP1* cells treated with IFNγ, Doxycycline (Dox) or left untreated and stained with CellMask. Mean fluorescence signal of the cytosol indicated by dashed red line and mean fluorescence intensity of the vacuoles (MFI) shown in table. N = number of vacuoles. **Data information:** *P* values from nested one-way ANOVA comparing means of n = 3 independent experiments from indicated condition to untreated WT cells.

**Fig. S2:**
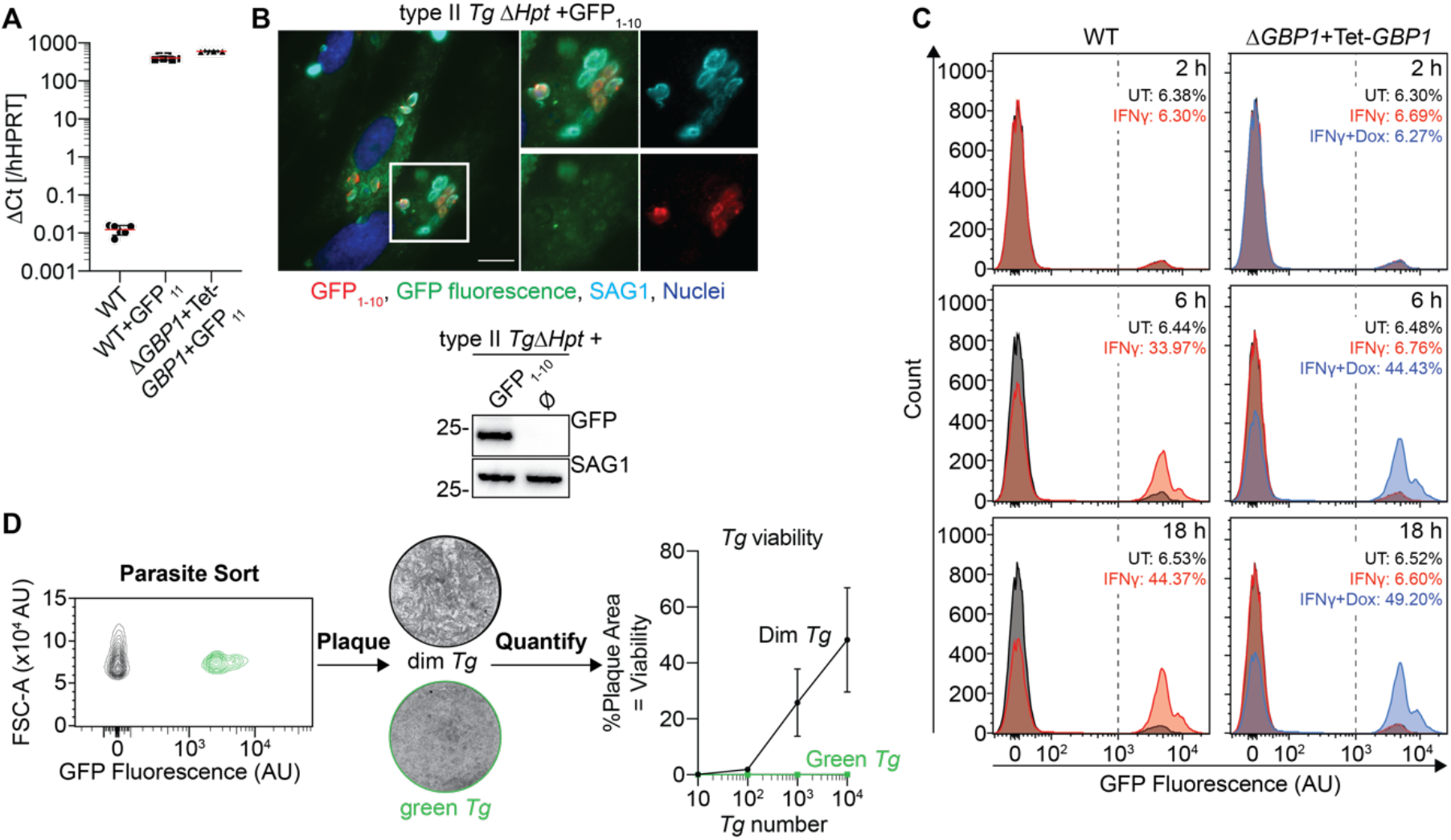
A novel split-GFP system allows quantitation of *Toxoplasma* parasite disruption by GBP1. **(A)** RT-qPCR to confirm expression of GFP_11_ fragment in the cytosol of THP-1 WT and THP-1 Δ*GBP1*+Tet-*GBP1* cells transduced with GFP_11_-Lentiviral particles. **(B)** Top: Representative immunofluorescence images and Bottom: immunoblot from type II *Tg* Δ*Hpt*+GFP_1-10_ or type II *Tg* Δ*Hpt* to confirm expression of the GFP fragment and absence of GFP fluorescence. Red: anti-GFP for GFP_1-10_; Green: GFP fluorescence; White: *Tg* surface antigen 1 (SAG1); Blue: Nuclei. Scale bar 20 μm. **(C)** Representative flow cytometry analysis of proportion of GFP-fluorescing and thus disrupted parasites harvested from untreated (UT) or IFNγ-primed THP-1 WT+GFP_11_ or from untreated, IFNγ-only or IFNγ- and Doxycycline (Dox)-treated THP-1 Δ*GBP1*+Tet-*GBP1*+GFP_11_ cells at 2, 6 or 18 hours post infection. Proportion of fluorescing parasites above the threshold of 10^3^ AU indicated in the figure. **(D)** Viability determination of type II *Tg* Δ*HpT*+GFP_1-10_ parasites harvested from IFNγ-primed THP-1 WT+GFP_11_ cells at 18 hours p.i, sorted based on their fluorescence (left), plaqued onto HFF cells (middle) and quantification of plaque area depending on number of parasites used for plaque formation (right). **Data information:** Graphs show mean ± SEM in **(A)** from n = 6 and in **(D)** from n = 3 independent experiments.

**Fig. S3:**
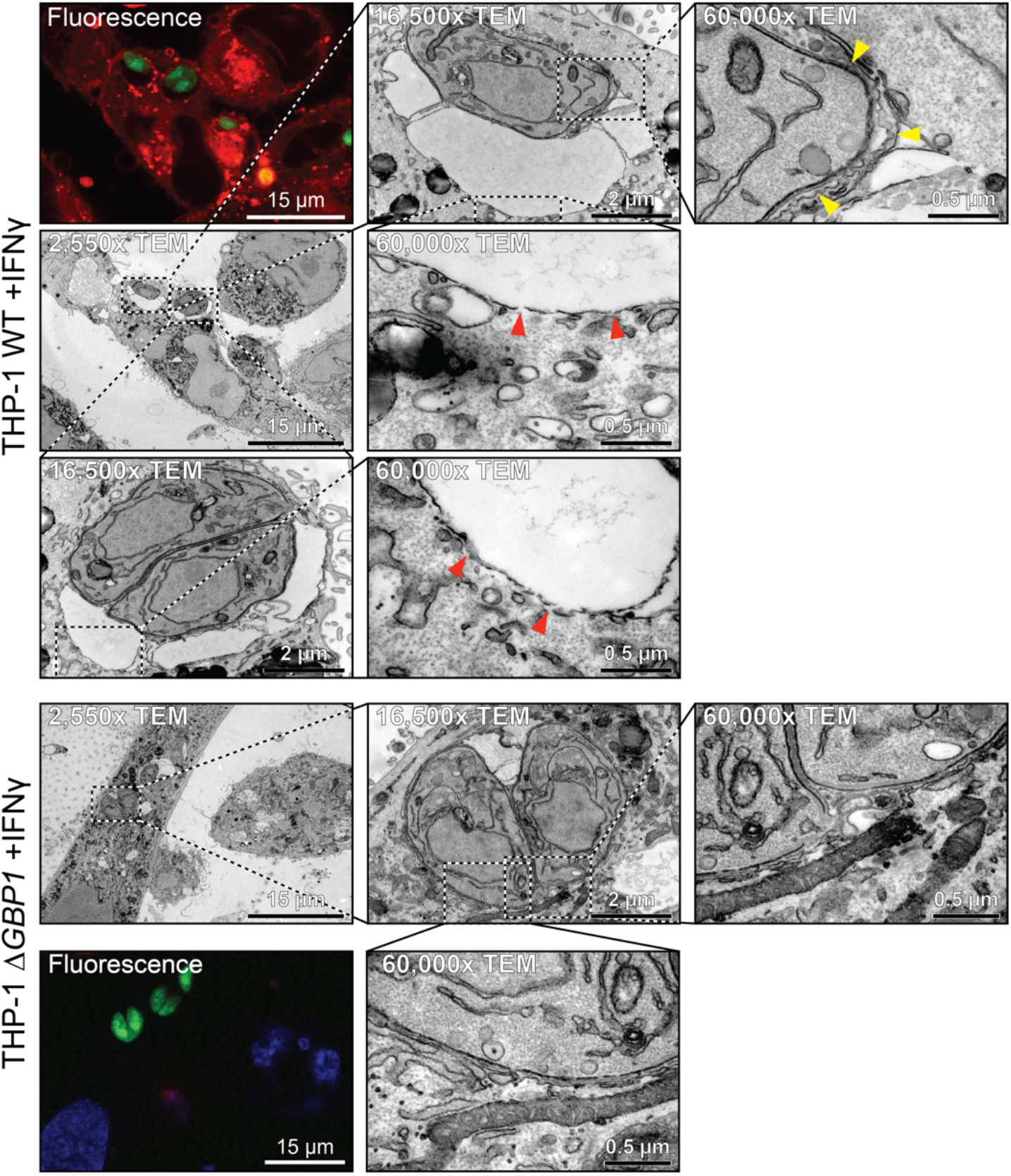
Correlative light and electron microscopy reveals ultrastructural defects of GBP1-targeted *Toxoplasma* vacuole membranes. Representative images of correlative light and electron microscopy of THP-1 WT or *ΔGBP1* cells (flooded with CellMask for fluorescence imaging), pre-treated with IFNγ to induce GBP1 expression and infected with type I (RH) *Toxoplasma gondii* (Tg) for 6 hours. Parasites indicated in boxes are shown at higher magnifications (TEM, transmission electron microscopy). Yellow arrowheads mark areas of vacuole membrane ruffling and red arrowheads mark areas of vacuole membrane disruption. Red: CellMask; Green: Tg; Blue: Nuclei. Scale bars as indicated.

**Fig. S4:**
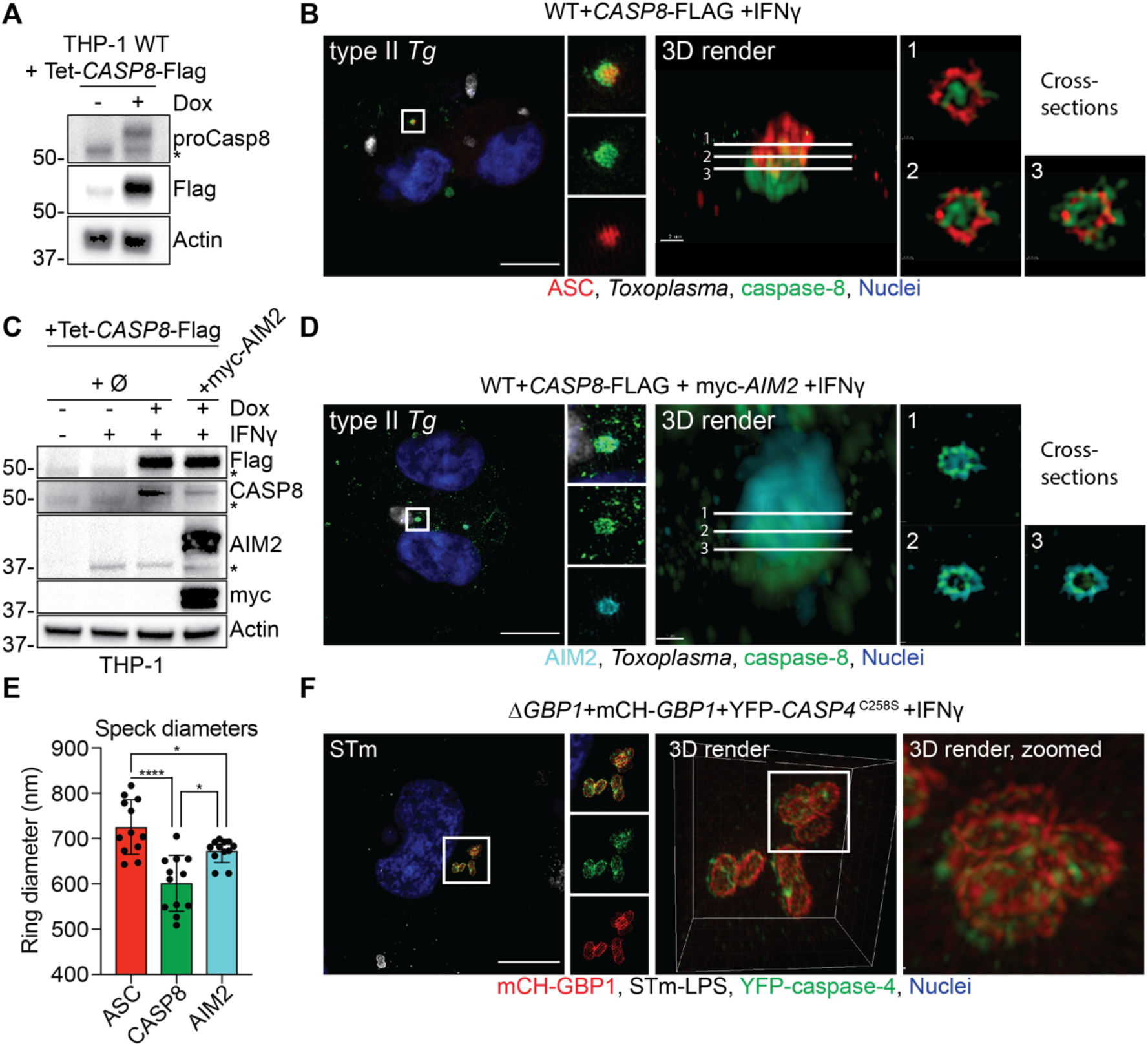
Structured illumination microscopy of caspase activation platforms formed by the action of GBP1. **(A)** Representative immunoblots for proCaspase-8, Flag and β-actin from THP-1 +Tet-*CASP8*-Flag cells showing Doxycycline (Dox)-inducible caspase-8-Flag expression. Cells were treated with Dox as indicated or left untreated. * endogenous caspase-8. **(B)** Left: Representative structured illumination immunofluorescence microscopy images from THP-1 WT+Tet-*CASP8*-Flag cells treated with IFNγ and Dox to induce caspase-8 expression and infected with type II *Toxoplasma gondii* (*Tg*) for 4 hours. Right: 3D reconstruction and slices through the ASC-caspase-8 speck. Red: ASC; Grey: *Tg*; Green: caspase-8; Blue: Nuclei. Scale bar 10 μm. **(C)** Representative immunoblots for Flag, caspase-8 (CASP8), AIM2, myc and β-actin from THP-1 WT, THP-1 +Tet-*CASP8*-Flag and THP-1 +Tet-*CASP8*-Flag+myc-*AIM2* cells showing Dox-inducible caspase-8-Flag expression and constitutive expression of myc-AIM2. Cells were treated with IFNγ, Dox or left untreated as indicated. * endogenous proteins. **(D)** Left: Representative structured illumination immunofluorescence microscopy images from THP-1+Tet-*CASP8*-Flag+myc-*AIM2* cells treated with IFNγ and Dox and infected with type II *Tg* for 4 hours. Right: 3D reconstruction and slices through the AIM2-caspase-8 speck. Cyan: AIM2; Grey: *Tg*; Green: caspase-8; Blue: Nuclei. Scale bar 10 μm. **(E)** Ring diameters of the indicated proteins within an inflammasome speck of cells shown in **(B)** and **(D)**. **(F)** Left: Representative structured illumination immunofluorescence microscopy images from THP-1 Δ*GBP1*+Tet-mCH-*GBP1*+YFP-*CASP4*^C258S^ cell treated with IFNγ and Dox and infected with *Salmonella* Typhimurium (STm) SL1344 (MOI = 30) for 2 hours. Right: 3D reconstruction of the GBP1-caspase-4 signalling platform on the cytosolic STm. Red: mCH-GBP1; Grey: STm-LPS; Green: YFP-caspase-4; Blue: Nuclei. Scale bar 10 μm. **Data information:** Graph in **(E)** shows quantification from n = 12 inflammasome specks and mean ± SEM. * *P* ≤ 0.05; **** *P* ≤ 0.0001 for indicated comparisons in **(E)** from one-way ANOVA following adjustment for multiple comparisons.

**Fig. S5:**
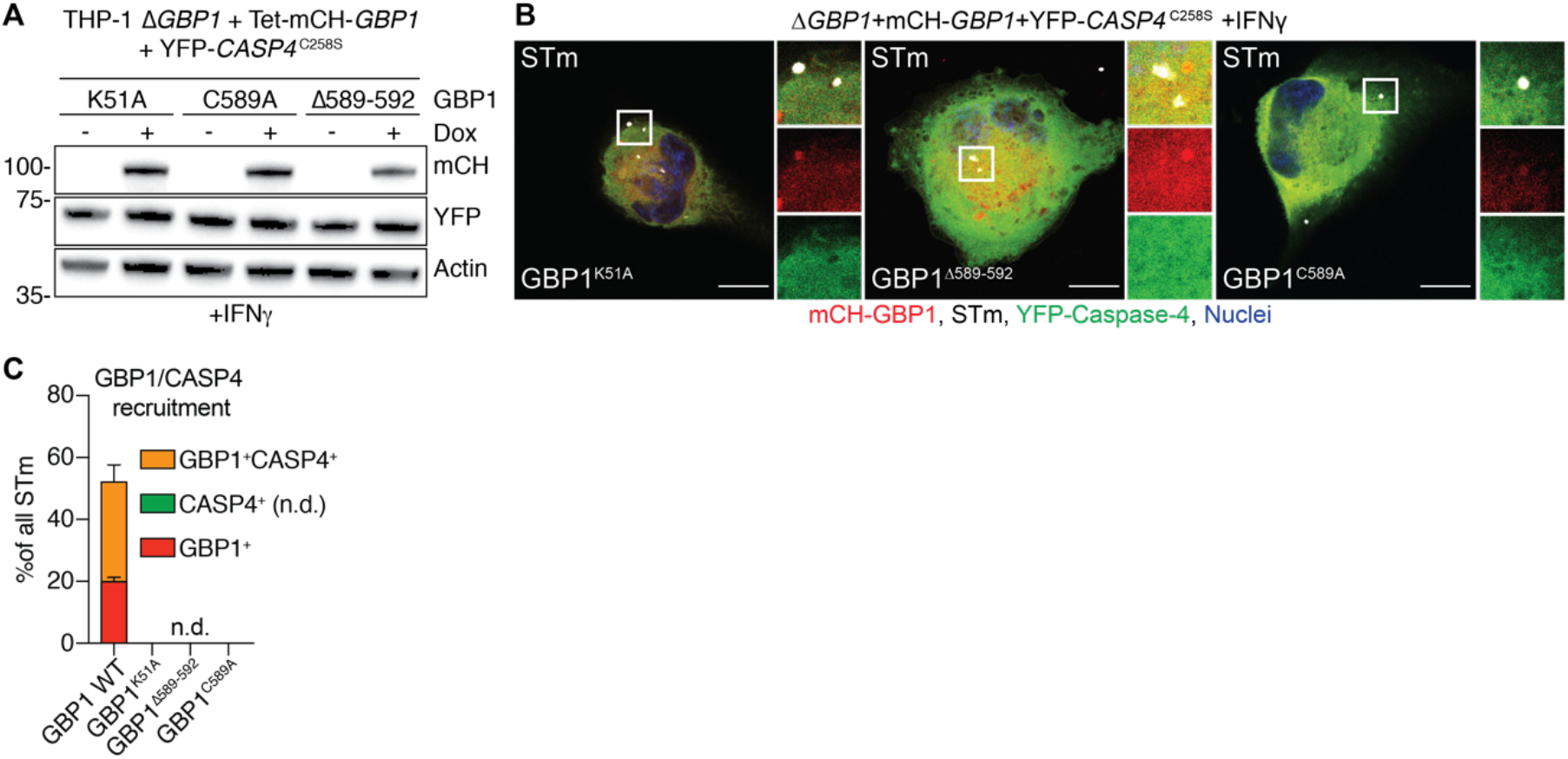
Molecular determinants of GBP1 and caspase-4 recruitment to cytosolic *Salmonella*. **(A)** Representative immunoblots for mCherry, YFP and β-actin from THP-1 Δ*GBP1*+Tet-mCH-*GBP1* cells expressing the indicated mutant of GBP1 and also stably expressing YFP-CASP4^C258S^. Cells were primed with IFNγ and treated with Doxycycline (Dox) as indicated. **(B)** Representative immunofluorescence images and **(C)** quantification of GBP1 and caspase-4 recruitment to *Salmonella* Typhimurium (STm) in IFNγ-primed and Dox-treated THP-1 Δ*GBP1*+Tet-mCH-*GBP1*+YFP-*CASP4*^C258S^ cells infected with STm SL1344 (MOI = 30) for 2 hours. Cells expressed the indicated mutants of GBP1. Red: mCH-GBP1; Grey: STm; Green: YFP-caspase-4; Blue: Nuclei. Scale bar 10 μm. **Data information:** Graph in **(C)** shows mean ± SEM of n = 3 independent experiments.

**Fig. S6:**
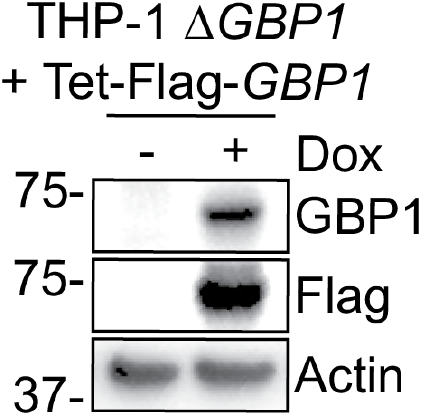
Verification immunoblot of THP-1 Δ*GBP1*+Tet-FLAG-*GBP1*. Representative immunoblots for GBP1, Flag and β-actin from THP-1 Δ*GBP1*+Tet-Flag-*GBP1* cells showing Dox-inducible Flag-GBP1 expression. Cells were treated with Doxycycline (Dox) as indicated or left untreated.

**Fig. S7:**
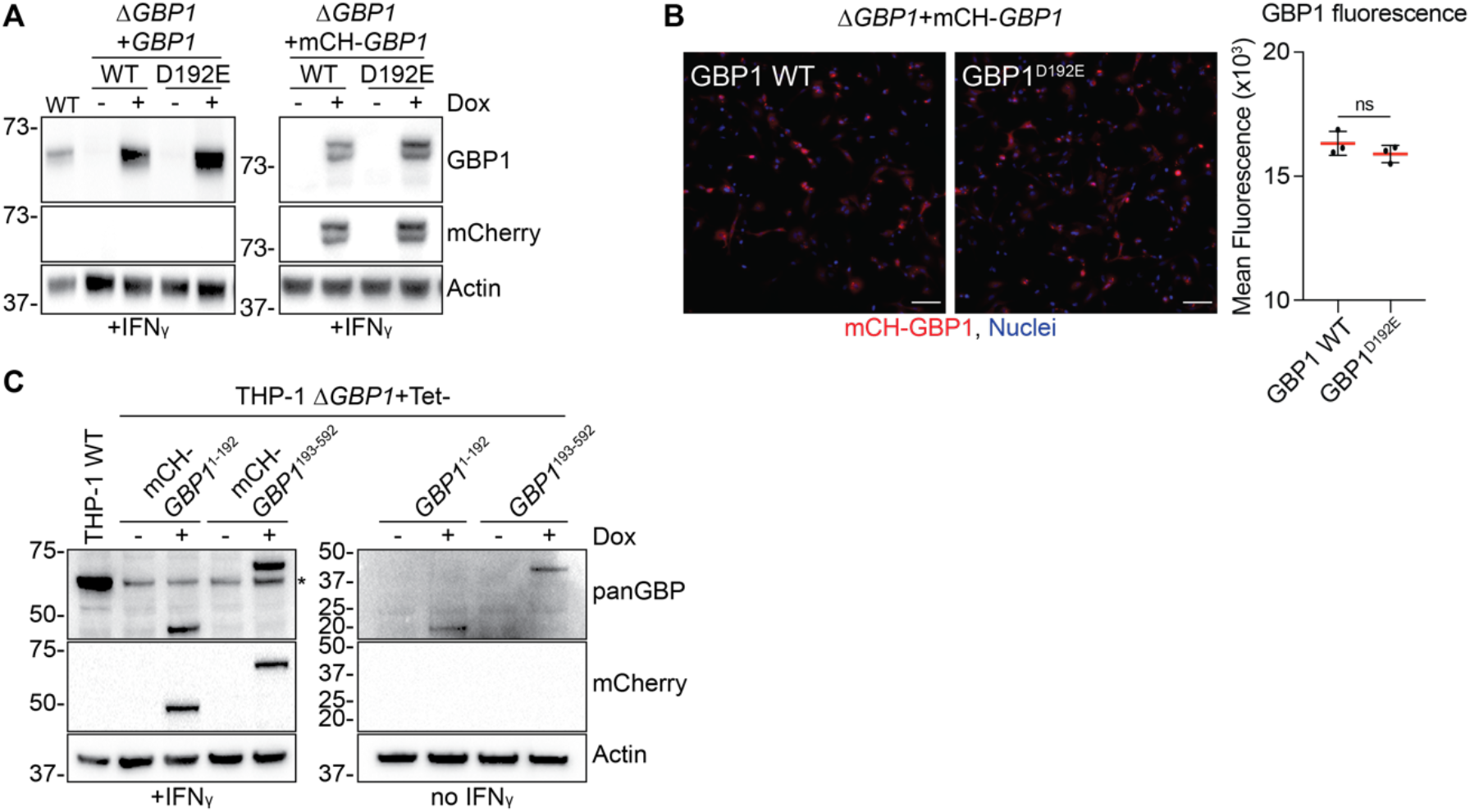
Verification and quality control of GBP1 D192 mutant and fragment cell lines. **(A)** Representative immunoblots of GBP1, mCherry (mCH) and β-actin from IFNγ-primed THP-1 Δ*GBP1*+Tet-*GBP1* or *GBP1*^D192E^ with and without mCH-tag for Doxycycline (Dox)-inducible expression. Cells were treated with Dox as indicated. **(B)** Representative images and quantification of mean fluorescence per cell from 100 fields of view from THP-1 Δ*GBP1*+Tet-mCH-*GBP1* or +Tet-mCH-*GBP1*^D192E^ cells treated with IFNγ and Dox to induce mCH-GBP1 expression. Red: mCH-GBP1; Blue: Nuclei. Scale bars 100 μm. **(C)** Immunoblot for human GBPs (panGBP), mCherry and Actin to confirm Dox-inducible expression of GBP1 fragments 1-192 or 193-592 with and without mCherry-tag in THP-1 Δ*GBP1*+Tet cells. * Marks endogenous, other GBP family members detected by the panGBP antibody in IFNγ-treated cells. **Data information:** Graph in **(B)** shows mean ± SEM from n = 3 independent experiments. *P* values in **(B)** from t-test. ns, not significant.

**Table S1.** Cell lines. Overview of cell lines used in this study.

| Cell line | Source |
| --- | --- |
| HEK293T | The Francis Crick Institute, Cell Services |
| HFF | The Francis Crick Institute, Cell Services |
| THP-1 ΔGBP1 | Fisch *et al.*, EMBOJ 2019 |
| THP-1 ΔGBP1 + Tet-EV | Fisch *et al.*, EMBOJ 2019 |
| THP-1 ΔGBP1 + Tet-GBP1 | Fisch *et al.*, EMBOJ 2019 |
| THP-1 ΔGBP1 + Tet-GBP1 + GFP<sub>11</sub> | Newly made cell line |
| THP-1 ΔGBP1 + Tet-GBP1<sup>1-192</sup> | Newly made cell line |
| THP-1 ΔGBP1 + Tet-GBP1<sup>193-592</sup> | Newly made cell line |
| THP-1 ΔGBP1 + Tet-GBP1<sup>D192E</sup> | Newly made cell line |
| THP-1 ΔGBP1 + Tet-mCH-GBP1 | Fisch *et al.*, EMBOJ 2019 |
| THP-1 ΔGBP1 + Tet-mCH-GBP1 + YFP-CASP4<sup>C258S</sup> | Fisch *et al.*, EMBOJ 2019 |
| THP-1 ΔGBP1 + Tet-mCH-GBP1<sup>Δ589-592</sup> + YFP-CASP4<sup>C258S</sup> | Newly made cell line |
| THP-1 ΔGBP1 + Tet-mCH-GBP1<sup>1-192</sup> | Newly made cell line |
| THP-1 ΔGBP1 + Tet-mCH-GBP1<sup>193-592</sup> | Newly made cell line |
| THP-1 ΔGBP1 + Tet-mCH-GBP1<sup>C589A</sup> + YFP-CASP4<sup>C258S</sup> | Newly made cell line |
| THP-1 ΔGBP1 + Tet-mCH-GBP1<sup>D192E</sup> | Newly made cell line |
| THP-1 ΔGBP1 + Tet-mCH-GBP1<sup>K51A</sup> + YFP-CASP4<sup>C258S</sup> | Newly made cell line |
| THP-1 WT | ATCC, TIB-202 |
| THP-1 WT + GFP<sub>11</sub> | Newly made cell line |
| THP-1 WT + Tet-CASP8 - Flag | Newly made cell line |
| THP-1 WT + Tet-CASP8 - Flag + myc-AIM2 | Newly made cell line |

**Table S2.** Antibodies. Overview of antibodies used in this study and indication of usage. Abbreviations: CST = Cell Signaling Technologies, IB = immunoblot, IF = immunofluorescence, IP = immunoprecipitation

| Antibody | Source | Use |
| --- | --- | --- |
| AIM2 | CST, #12948 | IB |
| ASC | Adipogen, AG-25B-0006 | IF |
| c-myc | Millipore, M2435 | IF, IB |
| CASP1 | Adipogen, AG-20B-0048 | IB |
| CASP8 | CST, #9746 | IB |
| Flag M2 | Millipore, F3165 | IF, IB, IP |
| Galectin-8 | R&D Systems, AF1305 | IF |
| GBP1 | Home-made (Frickel lab) | IB, IP |
| mCherry | Abcam, ab167453 | IB |
| panGBP | Home-made (Frickel lab) | IB |
| Rab7 | CST, #9367 | IF |
| STm | Abcam, ab35156 | IF |
| STm-LPS | Abcam, ab8274 | IF |
| YFP | Abcam, ab6556 | IB |
| $\beta$ -Actin | Sigma, A5316 | IB |

**Table S3.**
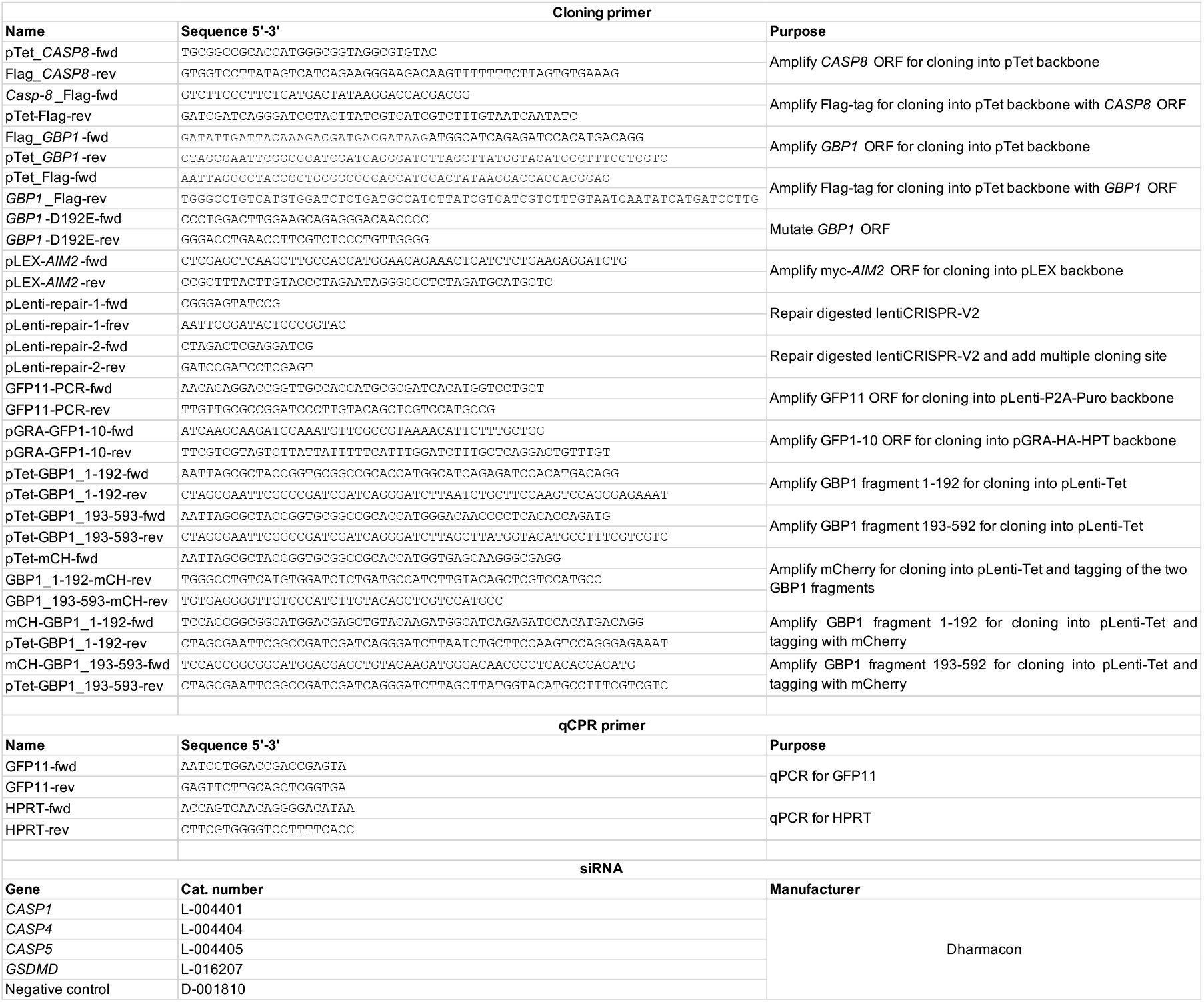
Primers and siRNAs. Overview of oligonucleotide primers and siRNAs used in this study.

**Table S4.** EM sample preparation protocol. Full BioWave program details for preparation of samples for electron microscopy.

| Description | Step# | Time<br>(min) | Time<br>(sec) | Power<br>(Watts) | SteadyTemp<br>temperature<br>(°C) | Vacuum cycle<br>vent time (sec) | Vacuum cycle<br>vacuum time<br>(sec) | Vacuum<br>set point<br>(inch Hg) | User Prompt<br>(1 = YES, 0 =<br>NO) | Vacuum OFF (1 =<br>no vacuum, 0 =<br>vacuum) | Vacuum<br>cycle (1 =<br>ON, 0 = OFF) | Vacuum ON (1 =<br>vacuum, 0 = no<br>vacuum) |
| --- | --- | --- | --- | --- | --- | --- | --- | --- | --- | --- | --- | --- |
| BENCH STEP Rinse in 0.1M PB | 1 | 0 | 0 | 0 | 21 | 0 | 0 | 0 | 1 | 1 | 0 | 0 |
| BENCH STEP Rinse in 0.1M PB | 2 | 0 | 0 | 0 | 21 | 0 | 0 | 0 | 1 | 1 | 0 | 0 |
| Rinse in 0.1M PB | 3 | 0 | 40 | 250 | 21 | 0 | 0 | 0 | 1 | 1 | 0 | 0 |
| Rinse in 0.1M PB | 4 | 0 | 40 | 250 | 21 | 0 | 0 | 0 | 1 | 1 | 0 | 0 |
| Osmium ON | 5 | 2 | 0 | 100 | 21 | 0 | 0 | 20 | 1 | 0 | 0 | 1 |
| Osmium OFF | 6 | 2 | 0 | 0 | 21 | 0 | 0 | 20 | 0 | 0 | 0 | 1 |
| Osmium ON | 7 | 2 | 0 | 100 | 21 | 0 | 0 | 20 | 0 | 0 | 0 | 1 |
| Osmium OFF | 8 | 2 | 0 | 0 | 21 | 0 | 0 | 20 | 0 | 0 | 0 | 1 |
| Osmium ON | 9 | 2 | 0 | 100 | 21 | 0 | 0 | 20 | 0 | 0 | 0 | 1 |
| Osmium OFF | 10 | 2 | 0 | 0 | 21 | 0 | 0 | 20 | 0 | 0 | 0 | 1 |
| Osmium ON | 11 | 2 | 0 | 100 | 21 | 0 | 0 | 20 | 0 | 0 | 0 | 1 |
| BENCH STEP Rinse in water | 12 | 0 | 0 | 0 | 21 | 0 | 0 | 0 | 1 | 1 | 0 | 0 |
| BENCH STEP Rinse in water | 13 | 0 | 0 | 0 | 21 | 0 | 0 | 0 | 1 | 1 | 0 | 0 |
| Rinse in water | 14 | 0 | 40 | 250 | 21 | 0 | 0 | 0 | 1 | 1 | 0 | 0 |
| Rinse in water | 15 | 0 | 40 | 250 | 21 | 0 | 0 | 0 | 1 | 1 | 0 | 0 |
| Thiocarbohydrazide ON | 16 | 2 | 0 | 100 | 40 | 0 | 0 | 20 | 1 | 0 | 0 | 1 |
| Thiocarbohydrazide OFF | 17 | 2 | 0 | 0 | 40 | 0 | 0 | 20 | 0 | 0 | 0 | 1 |
| Thiocarbohydrazide ON | 18 | 2 | 0 | 100 | 40 | 0 | 0 | 20 | 0 | 0 | 0 | 1 |
| Thiocarbohydrazide OFF | 19 | 2 | 0 | 0 | 40 | 0 | 0 | 20 | 0 | 0 | 0 | 1 |
| Thiocarbohydrazide ON | 20 | 2 | 0 | 100 | 40 | 0 | 0 | 20 | 0 | 0 | 0 | 1 |
| Thiocarbohydrazide OFF | 21 | 2 | 0 | 0 | 40 | 0 | 0 | 20 | 0 | 0 | 0 | 1 |
| Thiocarbohydrazide ON | 22 | 2 | 0 | 100 | 40 | 0 | 0 | 20 | 0 | 0 | 0 | 1 |
| BENCH STEP Rinse in water | 23 | 0 | 0 | 0 | 21 | 0 | 0 | 0 | 1 | 1 | 0 | 0 |
| BENCH STEP Rinse in water | 24 | 0 | 0 | 0 | 21 | 0 | 0 | 0 | 1 | 1 | 0 | 0 |
| Rinse in water | 25 | 0 | 40 | 250 | 21 | 0 | 0 | 0 | 1 | 1 | 0 | 0 |
| Rinse in water | 26 | 0 | 40 | 250 | 21 | 0 | 0 | 0 | 1 | 1 | 0 | 0 |
| Osmium ON | 27 | 2 | 0 | 100 | 21 | 0 | 0 | 20 | 1 | 0 | 0 | 1 |
| Osmium OFF | 28 | 2 | 0 | 0 | 21 | 0 | 0 | 20 | 0 | 0 | 0 | 1 |
| Osmium ON | 29 | 2 | 0 | 100 | 21 | 0 | 0 | 20 | 0 | 0 | 0 | 1 |
| Osmium OFF | 30 | 2 | 0 | 0 | 21 | 0 | 0 | 20 | 0 | 0 | 0 | 1 |
| Osmium ON | 31 | 2 | 0 | 100 | 21 | 0 | 0 | 20 | 0 | 0 | 0 | 1 |
| Osmium OFF | 32 | 2 | 0 | 0 | 21 | 0 | 0 | 20 | 0 | 0 | 0 | 1 |
| Osmium ON | 33 | 2 | 0 | 100 | 21 | 0 | 0 | 20 | 0 | 0 | 0 | 1 |
| BENCH STEP Rinse in water | 34 | 0 | 0 | 0 | 21 | 0 | 0 | 0 | 1 | 1 | 0 | 0 |
| BENCH STEP Rinse in water | 35 | 0 | 0 | 0 | 21 | 0 | 0 | 0 | 1 | 1 | 0 | 0 |
| Rinse in water | 36 | 0 | 40 | 250 | 21 | 0 | 0 | 0 | 1 | 1 | 0 | 0 |
| Rinse in water | 37 | 0 | 40 | 250 | 21 | 0 | 0 | 0 | 1 | 1 | 0 | 0 |
| Uranyl acetate ON | 38 | 2 | 0 | 100 | 40 | 0 | 0 | 20 | 1 | 0 | 0 | 1 |
| Uranyl acetate OFF | 39 | 2 | 0 | 0 | 40 | 0 | 0 | 20 | 0 | 0 | 0 | 1 |
| Uranyl acetate ON | 40 | 2 | 0 | 100 | 40 | 0 | 0 | 20 | 0 | 0 | 0 | 1 |
| Uranyl acetate OFF | 41 | 2 | 0 | 0 | 40 | 0 | 0 | 20 | 0 | 0 | 0 | 1 |
| Uranyl acetate ON | 42 | 2 | 0 | 100 | 40 | 0 | 0 | 20 | 0 | 0 | 0 | 1 |
| Uranyl acetate OFF | 43 | 2 | 0 | 0 | 40 | 0 | 0 | 20 | 0 | 0 | 0 | 1 |
| Uranyl acetate ON | 44 | 2 | 0 | 100 | 40 | 0 | 0 | 20 | 0 | 0 | 0 | 1 |
| BENCH STEP Rinse in water | 45 | 0 | 0 | 0 | 40 | 0 | 0 | 0 | 1 | 1 | 0 | 0 |
| BENCH STEP Rinse in water | 46 | 0 | 0 | 0 | 40 | 0 | 0 | 0 | 1 | 1 | 0 | 0 |
| Rinse in water | 47 | 0 | 45 | 250 | 40 | 0 | 0 | 0 | 1 | 1 | 0 | 0 |
| Rinse in water | 48 | 0 | 45 | 250 | 40 | 0 | 0 | 0 | 1 | 1 | 0 | 0 |
| Lead aspartate ON | 49 | 2 | 0 | 100 | 50 | 0 | 0 | 20 | 1 | 0 | 0 | 1 |
| Lead aspartate OFF | 50 | 2 | 0 | 0 | 50 | 0 | 0 | 20 | 0 | 0 | 0 | 1 |
| Lead aspartate ON | 51 | 2 | 0 | 100 | 50 | 0 | 0 | 20 | 0 | 0 | 0 | 1 |
| Lead aspartate OFF | 52 | 2 | 0 | 0 | 50 | 0 | 0 | 20 | 0 | 0 | 0 | 1 |
| Lead aspartate ON | 53 | 2 | 0 | 100 | 50 | 0 | 0 | 20 | 0 | 0 | 0 | 1 |
| Lead aspartate OFF | 54 | 2 | 0 | 0 | 50 | 0 | 0 | 20 | 0 | 0 | 0 | 1 |
| Lead aspartate ON | 55 | 2 | 0 | 100 | 50 | 0 | 0 | 20 | 0 | 0 | 0 | 1 |
| BENCH STEP Rinse in water | 56 | 0 | 0 | 0 | 21 | 0 | 0 | 0 | 1 | 1 | 0 | 0 |
| BENCH STEP Rinse in water | 57 | 0 | 0 | 0 | 21 | 0 | 0 | 0 | 1 | 1 | 0 | 0 |
| Rinse in water | 58 | 0 | 45 | 250 | 21 | 0 | 0 | 0 | 1 | 1 | 0 | 0 |
| Rinse in water | 59 | 0 | 45 | 250 | 21 | 0 | 0 | 0 | 1 | 1 | 0 | 0 |
| 70% Ethanol ON | 60 | 0 | 40 | 250 | 21 | 0 | 0 | 0 | 1 | 1 | 0 | 0 |
| 70% Ethanol ON | 61 | 0 | 40 | 250 | 21 | 0 | 0 | 0 | 1 | 1 | 0 | 0 |
| 90% Ethanol ON | 62 | 0 | 40 | 250 | 21 | 0 | 0 | 0 | 1 | 1 | 0 | 0 |
| 90% Ethanol ON | 63 | 0 | 40 | 250 | 21 | 0 | 0 | 0 | 1 | 1 | 0 | 0 |
| 100% Ethanol ON | 64 | 0 | 40 | 250 | 21 | 0 | 0 | 0 | 1 | 1 | 0 | 0 |
| 100% Ethanol ON | 65 | 0 | 40 | 250 | 21 | 0 | 0 | 0 | 1 | 1 | 0 | 0 |
| 50% Resin ON | 66 | 3 | 0 | 250 | 21 | 30 | 30 | 20 | 1 | 0 | 1 | 0 |
| 100% Resin ON | 67 | 3 | 0 | 250 | 21 | 30 | 30 | 20 | 1 | 0 | 1 | 0 |
| 100% Resin ON | 68 | 3 | 0 | 250 | 21 | 30 | 30 | 20 | 1 | 0 | 1 | 0 |
| 100% Resin ON | 69 | 3 | 0 | 250 | 21 | 30 | 30 | 20 | 1 | 0 | 1 | 0 |
| 100% Resin ON | 70 | 3 | 0 | 250 | 21 | 30 | 30 | 20 | 1 | 0 | 1 | 0 |
| TURN SYSTEM OFF | 71 | 0 | 0 | 0 | 21 | 0 | 0 | 0 | 0 | 1 | 0 | 0 |

## REFERENCES

Al-Zeer MA, Al-Younes HM, Braun PR, Zerrahn J & Meyer TF (2009) IFN-γ-Inducible Irga6 Mediates Host Resistance against Chlamydia trachomatis via Autophagy. PLoS One 4: e4588

Al-zeer MA, Al-younes HM, Lauster D, Lubad MA & Meyer TF (2013) Autophagy restricts Chlamydia trachomatis growth in human macrophages via IFNG-inducible guanylate binding proteins. Autophagy 9: 50-62

de Almeida L, Khare S, Misharin A V., Patel R, Ratsimandresy RA, Wallin MC, Perlman H, Greaves DR, Hoffman HM, Dorfleutner A & Stehlik C (2015) The PYRIN domain-only protein POP1 inhibits inflammasome assembly and ameliorates inflammatory disease. Immunity 43: 264

Balakrishnan A, Karki R, Berwin B, Yamamoto M & Kanneganti T-D (2018) Guanylate binding proteins facilitate caspase-11-dependent pyroptosis in response to type 3 secretion system-negative Pseudomonas aeruginosa. Cell Death Discov. 4: 66

Barz B, Loschwitz J & Strodel B (2019) Large-scale, dynamin-like motions of the human guanylate binding protein 1 revealed by multiresolution simulations. PLoS Comput. Biol. 15: e1007193

Bekpen C, Hunn JP, Rohde C, Parvanova I, Guethlein L, Dunn DM, Glowalla E, Leptin M & Howard JC (2005) The interferon-inducible p47 (IRG) GTPases in vertebrates: loss of the cell autonomous resistance mechanism in the human lineage. Genome Biol. 6: R92

Bekpen C, Xavier RJ & Eichler EE (2010) Human IRGM gene “to be or not to be.” Semin. Immunopathol. 32: 437–444

Bernstein-Hanley I, Coers J, Balsara ZR, Taylor GA, Starnbach MN & Dietrich WF (2006) The p47 GTPases Igtp and Irgb10 map to the Chlamydia trachomatis susceptibility locus Ctrq-3 and mediate cellular resistance in mice. Proc. Natl. Acad. Sci. U.S.A. 103: 14092–14097

Bradfield C (2016) Sulfated DAMPs mobilize human GBPs for cell-autonomous immunity against bacterial pathogens.

Brest P, Lapaquette P, Souidi M, Lebrigand K, Cesaro A, Vouret-Craviari V, Mari B, Barbry P, Mosnier JF, Hébuterne X, Harel-Bellan A, Mograbi B, Darfeuille-Michaud A & Hofman P (2011) A synonymous variant in IRGM alters a binding site for miR-196 and causes deregulation of IRGM-dependent xenophagy in Crohn’s disease. Nat. Genet. 43: 242–245

Britzen-Laurent N, Bauer M, Berton V, Fischer N, Syguda A, Reipschläger S, Naschberger E, Herrmann C & Stürzl M (2010) Intracellular trafficking of guanylate-binding proteins is regulated by heterodimerization in a hierarchical manner. PLoS One 5: e14246

Burleigh K, Maltbaek JH, Cambier S, Green R, Gale M, James RC & Stetson DB (2020) Human DNA-PK activates a STING-independent DNA sensing pathway. Sci. Immunol. 5: eaba4219

Cardona A, Saalfeld S, Schindelin J, Arganda-Carreras I, Preibisch S, Longair M, Tomancak P, Hartenstein V & Douglas RJ (2012) TrakEM2 Software for Neural Circuit Reconstruction. PLoS One 7: e38011

Casson CN, Yu J, Reyes VM, Taschuk FO, Yadav A, Copenhaver AM, Nguyen HT, Collman RG & Shin S (2015) Human caspase-4 mediates noncanonical inflammasome activation against gram negative bacterial pathogens. Proc. Natl. Acad. Sci. U.S.A. 112: 6688–6693

Cheng YS, Patterson CE & Staeheli P (1991) Interferon-induced guanylate-binding proteins lack an N(T)KXD consensus motif and bind GMP in addition to GDP and GTP. Mol. Cell. Biol. 11: 4717–4725

Coers J, Bernstein-Hanley I, Grotsky D, Parvanova I, Howard JC, Taylor GA, Dietrich WF & Starnbach MN (2008) Chlamydia muridarum Evades Growth Restriction by the IFN-γ-Inducible Host Resistance Factor Irgb10. J. Immunol. 180: 6237–6245

Coppens I, Dunn JD, Romano JD, Pypaert M, Zhang H, Boothroyd JC & Joiner KA (2006) Toxoplasma gondii Sequesters Lysosomes from Mammalian Hosts in the Vacuolar Space. Cell 125: 261–274

Costa Franco MM, Marim F, Guimarães ES, Assis NRG, Cerqueira DM, Alves-Silva J, Harms J, Splitter G, Smith J, Kanneganti T-D, de Queiroz NMGP, Gutman D, Barber GN & Oliveira SC (2018) Brucella abortus Triggers a cGAS-Independent STING Pathway To Induce Host Protection That Involves Guanylate-Binding Proteins and Inflammasome Activation. J. Immunol. 200: 607–622

Creasey EA & Isberg RR (2012) The protein SdhA maintains the integrity of the Legionella-containing vacuole. Proc. Natl. Acad. Sci. U. S. A. 109: 3481–3486

Deerinck TJ, Bushong E, Thor A, Ellisman M, Deerinck T, Thor A, Deerinck T, Bushong E, Thor CA & Ellisman M (2010) NCMIR methods for 3D EM: A new protocol for preparation of biological specimens for serial block face scanning electron microscopy. Microscopy

Degrandi D, Konermann C, Beuter-Gunia C, Kresse A, Würthner J, Kurig S, Beer S & Pfeffer K (2007) Extensive Characterization of IFN-Induced GTPases mGBP1 to mGBP10 Involved in Host Defense. J. Immunol. 179: 7729–7740

Degrandi D, Kravets E, Konermann C, Beuter-Gunia C, Klumpers V, Lahme S, Wischmann E, Mausberg AK, Beer-Hammer S, Pfeffer K, Klümpers V, Lahme S, Wischmann E, Mausberg AK, Beer-Hammer S & Pfeffer K (2013) Murine Guanylate Binding Protein 2 (mGBP2) controls Toxoplasma gondii replication. Proc. Natl. Acad. Sci. U.S.A. 110: 294–9

Ding J & Shao F (2017) SnapShot: The Noncanonical Inflammasome. Cell 168: 544–544.e1

Eldridge MJG, Sanchez-Garrido J, Hoben GF, Goddard PJ & Shenoy AR (2017) The Atypical Ubiquitin E2 Conjugase UBE2L3 Is an Indirect Caspase-1 Target and Controls IL-1β Secretion by Inflammasomes. Cell Rep. 18: 1285–1297

Feeley EM, Pilla-Moffett DM, Zwack EE, Piro AS, Finethy R, Kolb JP, Martinez J, Brodsky IE & Coers J (2017) Galectin-3 directs antimicrobial guanylate binding proteins to vacuoles furnished with bacterial secretion systems. Proc. Natl. Acad. Sci. U.S.A. 114: E1698–E1706

Fisch D, Bando H, Clough B, Hornung V, Yamamoto M, Shenoy AR & Frickel E (2019a) Human GBP1 is a microbe-specific gatekeeper of macrophage apoptosis and pyroptosis. EMBO J. 38: e100926

Fisch D, Yakimovich A, Clough B, Wright J, Bunyan M, Howell M, Mercer J & Frickel E (2019b) Defining host-pathogen interactions employing an artificial intelligence workflow. Elife 8: e40560

Foltz C, Napolitano A, Khan R, Clough B, Hirst EM & Frickel E-M (2017) TRIM21 is critical for survival of Toxoplasma gondii infection and localises to GBP-positive parasite vacuoles. Sci. Rep. 7: 5209

Forster F, Paster W, Supper V, Schatzlmaier P, Sunzenauer S, Ostler N, Saliba A, Eckerstorfer P, Britzen-Laurent N, Schutz G, Schmid JA, Zlabinger GJ, Naschberger E, Sturzl M & Stockinger H (2014) Guanylate Binding Protein 1-Mediated Interaction of T Cell Antigen Receptor Signaling with the Cytoskeleton. J. Immunol. 192: 771–781

Fres JM, Muller S & Praefcke GJK (2010) Purification of the CaaX-modified, dynamin-related large GTPase hGBP1 by coexpression with farnesyltransferase. J. Lipid Res. 51: 2454–2459

Ghosh A, Praefcke GJKK, Renault L, Wittinghofer A & Herrmann C (2006) How guanylate-binding proteins achieve assembly-stimulated processive cleavage of GTP to GMP. Nature 440: 101–104

Gomes MTR, Cerqueira DM, Guimarães ES, Campos PC & Oliveira SC (2019) Guanylate-binding proteins at the crossroad of noncanonical inflammasome activation during bacterial infections. J. Leukoc. Biol. 106: 553–562

Hagar JA, Powell DA, Aachoui Y, Ernst RK & Miao EA (2013) Cytoplasmic LPS activates caspase-11: Implications in TLR4-independent endotoxic shock. Science 341: 1250–1253

Haldar AK, Foltz C, Finethy R, Piro AS, Feeley EM, Pilla-Moffett DM, Komatsu M, Frickel E-M & Coers J (2015) Ubiquitin systems mark pathogen-containing vacuoles as targets for host defense by guanylate binding proteins. Proc. Natl. Acad. Sci. U.S.A. 112: E5628–E5637

Haldar AK, Piro AS, Pilla DM, Yamamoto M & Coers J (2014) The E2-like conjugation enzyme Atg3 promotes binding of IRG and Gbp proteins to Chlamydia- and Toxoplasma-containing vacuoles and host resistance. PLoS One 9: e86684

Haldar AK, Saka HA, Piro AS, Dunn JD, Henry SC, Taylor GA, Frickel EM, Valdivia RH & Coers J (2013) IRG and GBP host resistance factors target aberrant, non-self vacuoles characterized by the missing of self IRGM proteins. PLoS Pathog. 9: e1003414

Henry SC, Daniell X, Indaram M, Whitesides JF, Sempowski GD, Howell D, Oliver T & Taylor GA (2007) Impaired Macrophage Function Underscores Susceptibility to Salmonella in Mice Lacking Irgm1 (LRG-47). J. Immunol. 179: 6963–6972

Huang S, Meng Q, Maminska A & MacMicking JD (2019) Cell-autonomous immunity by IFN-induced GBPs in animals and plants. Curr. Opin. Immunol. 60: 71–80

Hunn JP, Koenen-Waisman S, Papic N, Schroeder N, Pawlowski N, Lange R, Kaiser F, Zerrahn J, Martens S & Howard JC (2008) Regulatory interactions between IRG resistance GTPases in the cellular response to Toxoplasma gondii. EMBO J. 27: 2495–2509

Ince S, Kutsch M, Shydlovskyi S & Herrmann C (2017) The human guanylate-binding proteins hGBP-1 and hGBP-5 cycle between monomers and dimers only. FEBS J. 284: 2284–2301

Johnston AC, Piro A, Clough B, Siew M, Virreira Winter S, Coers J & Frickel E-M (2016) Human GBP1 does not localize to pathogen vacuoles but restricts Toxoplasma gondii. Cell. Microbiol. 18: 1056–64

Jorgensen I, Rayamajhi M & Miao EA (2017) Programmed cell death as a defence against infection. Nat. Rev. Immunol. 17: 151–164

Jorgensen I, Zhang Y, Krantz BA & Miao EA (2016) Pyroptosis triggers pore-induced intracellular traps (PITs) that capture bacteria and lead to their clearance by efferocytosis. J. Exp. Med. 213: 2113–2128

Kagan JC, Magupalli VG & Wu H (2014) SMOCs: supramolecular organizing centres that control innate immunity. Nat. Rev. Immunol. 14: 821

Kayagaki N, Warming S, Lamkanfi M, Walle L Vande, Louie S, Dong J, Newton K, Qu Y, Liu J, Heldens S, Zhang J, Lee WP, Roose-Girma M & Dixit VM (2011) Non-canonical inflammasome activation targets caspase-11. Nature 479: 117–121

Kayagaki N, Wong MT, Stowe IB, Ramani SR, Gonzalez LC, Akashi-Takamura S, Miyake K, Zhang J, Lee WP, Muszynśki A, Forsberg LS, Carlson RW & Dixit VM (2013) Noncanonical inflammasome activation by intracellular LPS independent of TLR4. Science 341: 1246–1249

Khaminets A, Hunn JP, Könen-Waisman S, Zhao YO, Preukschat D, Coers J, Boyle JP, Ong Y-CC, Boothroyd JC, Reichmann G & Howard JC (2010) Coordinated loading of IRG resistance GTPases on to the Toxoplasma gondii parasitophorous vacuole. Cell. Microbiol. 12: 939–961

Khare S, Ratsimandresy RA, de Almeida L, Cuda CM, Rellick SL, Misharin A V, Wallin MC, Gangopadhyay A, Forte E, Gottwein E, Perlman H, Reed JC, Greaves DR, Dorfleutner A & Stehlik C (2014) The PYRIN domain-only protein POP3 inhibits ALR inflammasomes and regulates responses to infection with DNA viruses. Nat. Immunol. 15: 343–53

Kim B-H, Shenoy AR, Kumar P, Bradfield CJ & MacMicking JD (2012a) IFN-Inducible GTPases in Host Cell Defense. Cell Host Microbe 12: 432–444

Kim B-H, Shenoy AR, Kumar P, Das R, Tiwari S & MacMicking JD (2011) A Family of IFN-γ-Inducible 65-kD GTPases Protects Against Bacterial Infection. Science 332: 717–721

Kim BH, Shenoy AR, Kumar P, Bradfield CJ & MacMicking JD (2012b) IFN-inducible GTPases in host cell defense. Cell Host Microbe 12: 432–444

Kohler KM, Kutsch M, Piro AS, Wallace G, Coers J & Barber MF (2019) A rapidly evolving polybasic motif modulates bacterial detection by guanylate binding proteins. bioRxiv: 689554

Kortmann J, Brubaker SW & Monack DM (2015) Cutting Edge: Inflammasome Activation in Primary Human Macrophages Is Dependent on Flagellin. J. Immunol. 195: 815–819

Kravets E, Degrandi D, Ma Q, Peulen T-OO, Klümpers V, Felekyan S, Kühnemuth R, Weidtkamp-Peters S, Seidel CA & Pfeffer K (2016) Guanylate binding proteins directly attack Toxoplasma gondii via supramolecular complexes. Elife 5: e11479

Kresse A, Konermann C, Degrandi D, Beuter-Gunia C, Wuerthner J, Pfeffer K, Beer S, Gupta S, Rubin B, Holmes S, Cheng Y, Colonno R, Yin F, Decker T, Lew D, Cheng Y, Levy D, Darnell J, Decker T, Lew D, et al (2008) Analyses of murine GBP homology clusters based on in silico, in vitro and in vivo studies. BMC Genomics 9: 158–170

Lagrange B, Benaoudia S, Wallet P, Magnotti F, Provost A, Michal F, Martin A, Di Lorenzo F, Py BF, Molinaro A & Henry T (2018) Human caspase-4 detects tetra-acylated LPS and cytosolic Francisella and functions differently from murine caspase-11. Nat. Commun. 9: 242

Lapaquette P, Glasser AL, Huett A, Xavier RJ & Darfeuille-Michaud A (2010) Crohn’s disease-associated adherent-invasive E. coli are selectively favoured by impaired autophagy to replicate intracellularly. Cell. Microbiol. 12: 99–113

Li P, Jiang W, Yu Q, Liu W, Zhou P, Li J, Xu J, Xu B, Wang F & Shao F (2017) Ubiquitination and degradation of GBPs by a Shigella effector to suppress host defence. Nature 551: 378

Lindenberg V, Mölleken K, Kravets E, Stallmann S, Hegemann JH, Degrandi D & Pfeffer K (2017) Broad recruitment of mGBP family members to Chlamydia trachomatis inclusions. PLoS One 12: e0185273

Liu BC, Sarhan J, Panda A, Muendlein HI, Ilyukha V, Coers J, Yamamoto M, Isberg RR & Poltorak A (2018) Constitutive Interferon Maintains GBP Expression Required for Release of Bacterial Components Upstream of Pyroptosis and Anti-DNA Responses. Cell Rep. 24: 155–168.e5

Lorenz C, Ince S, Zhang T, Cousin A, Batra-Safferling R, Nagel-Steger L, Herrmann C & Stadler AM (2019) Farnesylation of human guanylate binding protein 1 as safety mechanism preventing structural rearrangements and uninduced dimerization. FEBS J. 287: 496–514

MacMicking JD (2012) Interferon-inducible effector mechanisms in cell-autonomous immunity. Nat. Rev. Immunol. 12: 367–382

MacMicking JD, Taylor GA & McKinney JD (2003) Immune Control of Tuberculosis by IFN-γ-inducible LRG-47. Science 302: 654–659

Man SM, Hopkins LJ, Nugent E, Cox S, Glück IM, Tourlomousis P, Wright JA, Cicuta P, Monie TP & Bryant CE (2014) Inflammasome activation causes dual recruitment of NLRC4 and NLRP3 to the same macromolecular complex. Proc. Natl. Acad. Sci. U. S. A. 111: 7403–8

Man SM, Karki R, Malireddi RKS, Neale G, Vogel P, Yamamoto M, Lamkanfi M & Kanneganti T (2015) The transcription factor IRF1 and guanylate-binding proteins target activation of the AIM2 inflammasome by Francisella infection. Nat. Immunol. 16: 467–475

Man SM, Karki R, Sasai M, Place DE, Kesavardhana S, Temirov J, Frase S, Zhu Q, Malireddi RKS, Kuriakose T, Peters JL, Neale G, Brown SA, Yamamoto M & Kanneganti T-D (2016) IRGB10 Liberates Bacterial Ligands for Sensing by the AIM2 and Caspase-11-NLRP3 Inflammasomes. Cell: 1–15

Man SM, Place DE, Kuriakose T & Kanneganti T-D (2017) Interferon-inducible guanylate-binding proteins at the interface of cell-autonomous immunity and inflammasome activation. J. Leukoc. Biol. 101: 143–150

Man SM, Tourlomousis P, Hopkins L, Monie TP, Fitzgerald KA & Bryant CE (2013) Salmonella infection induces recruitment of Caspase-8 to the inflammasome to modulate IL-1β production. J. Immunol. 191: 5239–46

Meunier E & Broz P (2015) Quantification of Cytosolic vs. Vacuolar Salmonella in Primary Macrophages by Differential Permeabilization. J. Vis. Exp.: e52960

Meunier E & Broz P (2016) Interferon-inducible GTPases in cell autonomous and innate immunity. Cell. Microbiol. 18: 168–180

Meunier E, Dick MS, Dreier RF, Schürmann N, Broz DK, Warming S, Roose-Girma M, Bumann D, Kayagaki N, Takeda K, Yamamoto M & Broz P (2014) Caspase-11 activation requires lysis of pathogen-containing vacuoles by IFN-induced GTPases. Nature 509: 366–370

Meunier E, Wallet P, Dreier RF, Costanzo S, Anton L, Rühl S, Dussurgey S, Dick MS, Kistner A, Rigard M, Degrandi D, Pfeffer K, Yamamoto M, Henry T & Broz P (2015) Guanylate-binding proteins promote activation of the AIM2 inflammasome during infection with Francisella novicida. Nat. Immunol. 16: 476–84

Miyairi I, Tatireddigari VRRA, Mahdi OS, Rose LA, Belland RJ, Lu L, Williams RW & Byrne GI (2007) The p47 GTPases Iigp2 and Irgb10 Regulate Innate Immunity and Inflammation to Murine Chlamydia psittaci Infection. J. Immunol. 179: 1814–1824

Mostowy S & Shenoy AR (2015) The cytoskeleton in cell-autonomous immunity: structural determinants of host defence. Nat. Rev. Immunol. 15: 559–73

Nantais DE, Schwemmle M, Stickney JT, Vestal DJ & Buss JE (1996) Prenylation of an interferon-γ-induced GTP-binding protein: The human guanylate binding protein, huGBP1. J. Leukoc. Biol. 60: 423–431

Naschberger E, Geißdörfer W, Bogdan C, Tripal P, Kremmer E, Stürzl M & Britzen-Laurent N (2017) Processing and secretion of guanylate binding protein-1 depend on inflammatory caspase activity. J. Cell. Mol. Med. 20: 1–13

Olszewski MA, Gray J & Vestal DJ (2006) In Silico Genomic Analysis of the Human and Murine Guanylate-Binding Protein (GBP) Gene Clusters. J. Interf. Cytokine 352: 328–352

Ostler N, Britzen-Laurent N, Liebl A, Naschberger E, Lochnit G, Ostler M, Forster F, Kunzelmann P, Ince S, Supper V, Praefcke GJK, Schubert DW, Stockinger H, Herrmann C & Sturzl M (2014) Gamma Interferon-Induced Guanylate Binding Protein 1 Is a Novel Actin Cytoskeleton Remodeling Factor. Mol. Cell. Biol. 34: 196–209

Pilla-Moffett D, Barber MF, Taylor GA & Coers J (2016) Interferon-Inducible GTPases in Host Resistance, Inflammation and Disease. J. Mol. Biol. 428: 3495–3513

Pilla DM, Hagar JA, Haldar AK, Mason AK, Degrandi D, Pfeffer K, Ernst RK, Yamamoto M, Miao EA & Coers J (2014) Guanylate binding proteins promote caspase-11-dependent pyroptosis in response to cytoplasmic LPS. Proc. Natl. Acad. Sci. U.S.A. 111: 6046–51

Piro AS, Hernandez D, Luoma S, Feeley EM, Finethy R, Yirga A, Frickel EM, Lesser CF & Coers J (2017) Detection of Cytosolic Shigella flexneri via a C-Terminal Triple-Arginine Motif of GBP1 Inhibits Actin-Based Motility. MBio 8:

Praefcke GJK, Geyer M, Schwemmle M, Robert Kalbitzer H & Herrmann C (1999) Nucleotide-binding characteristics of human guanylate-binding protein 1 (hGBP1) and identification of the third GTP-binding motif. J. Mol. Biol. 292: 321–332

Prakash B, Praefcke GJK, Renault L, Wittinghofer A & Herrmann C (2000) Structure of human guanylate-binding protein 1 representing a unique class of GTP-binding proteins. Nature 403: 567–571

Randow F, MacMicking JD & James LC (2013) Cellular self-defense: how cell-autonomous immunity protects against pathogens. Science 340: 701–706

Reyes Ruiz VM, Ramirez J, Naseer N, Palacio NM, Siddarthan IJ, Yan BM, Boyer MA, Pensinger DA, Sauer J-D & Shin S (2017) Broad detection of bacterial type III secretion system and flagellin proteins by the human NAIP/NLRC4 inflammasome. Proc. Natl. Acad. Sci. U. S. A. 114: 13242–13247

Romei MG & Boxer SG (2019) Split Green Fluorescent Proteins: Scope, Limitations, and Outlook. Annu. Rev. Biophys. 48: 19–44

Rühl S & Broz P (2015) Caspase-11 activates a canonical NLRP3 inflammasome by promoting K+ efflux. Eur. J. Immunol. 45: 2927–2936

Saeij JP & Frickel E-M (2017) Exposing Toxoplasma gondii hiding inside the vacuole: a role for GBPs, autophagy and host cell death. Curr. Opin. Microbiol. 40: 72–80

Sanchez-Garrido J, Sancho-Shimizu V & Shenoy AR (2018) Regulated proteolysis of p62/SQSTM1 enables differential control of autophagy and nutrient sensing. Sci. Signal. 11: eaat6903

Sanjana NE, Shalem O & Zhang F (2014) Improved vectors and genome-wide libraries for CRISPR screening. Nat. Methods 11: 783–784

Santos JC, Dick MS, Lagrange B, Degrandi D, Pfeffer K, Yamamoto M, Meunier E, Pelczar P, Henry T & Broz P (2018) LPS targets host guanylate-binding proteins to the bacterial outer membrane for non-canonical inflammasome activation. EMBO J. 37:

Schoggins JW (2019) Interferon-Stimulated Genes: What Do They All Do? Annu. Rev. Virol. 18:

Schwemmlel M & Staeheli P (1994) The interferon-induced 67-kDa guanylate-binding protein (hGBP1) is a GTPase that converts GTP to GMP. J. Biol. Chem. 269: 11299–11305

Selleck EM, Fentress SJ, Beatty WL, Degrandi D, Pfeffer K, Virgin HW, MacMicking JD & Sibley LD (2013) Guanylate-binding Protein 1 (Gbp1) Contributes to Cell-autonomous Immunity against Toxoplasma gondii. PLoS Pathog. 9: e1003320

Shenoy AR, Kim BH, Choi HP, Matsuzawa T, Tiwari S & MacMicking JD (2007) Emerging themes in IFN-γ-induced macrophage immunity by the p47 and p65 GTPase families. Immunobiology 212: 771–784

Shenoy AR, Wellington DA, Kumar P, Kassa H, Booth CJ, Cresswell P & MacMicking JD (2012) GBP5 Promotes NLRP3 Inflammasome Assembly and Immunity in Mammals. Science 336: 481–485

Shi J, Zhao Y, Wang Y, Gao W, Ding J, Li P, Hu L & Shao F (2014) Inflammatory caspases are innate immune receptors for intracellular LPS. Nature 514: 187–192

Shydlovskyi S, Zienert AY, Ince S, Dovengerds C, Hohendahl A, Dargazanli JM, Blum A, Günther SD, Kladt N, Stürzl M, Schauss AC, Kutsch M, Roux A, Praefcke GJK & Herrmann C (2017) Nucleotide-dependent farnesyl switch orchestrates polymerization and membrane binding of human guanylate-binding protein 1. Proc. Natl. Acad. Sci. U. S. A. 114: E5559–E5568

Singh SB, Davis AS, Taylor GA & Deretic V (2006) Human IRGM induces autophagy to eliminate intracellular mycobacteria. Science 313: 1438–1441

Singh SB, Ornatowski W, Vergne I, Naylor J, Delgado M, Roberts E, Ponpuak M, Master S, Pilli M, White E, Komatsu M & Deretic V (2010) Human IRGM regulates autophagy and cell-autonomous immunity functions through mitochondria. Nat. Cell Biol. 12: 1154–1165

Stennicke HR & Salvesen GS (1997) Biochemical characteristics of caspases-3, −6, −7, and −8. J. Biol. Chem. 272: 25719–23

Stickney JT & Buss JE (2000) Murine guanylate-binding protein: Incomplete geranylgeranyl isoprenoid modification of an interferon-γ-inducible guanosine triphosphate binding protein. Mol. Biol. Cell 11: 2191–2200

Thurston TLMM, Matthews SA, Jennings E, Alix E, Shao F, Shenoy AR, Birrell MA & Holden DW (2016) Growth inhibition of cytosolic Salmonella by caspase-1 and caspase-11 precedes host cell death. Nat. Commun. 7: 13292

Thurston TLMM, Wandel MP, Von Muhlinen N, Foeglein Á, Randow F, Foeglein A, Randow F, Foeglein Á & Randow F (2012) Galectin 8 targets damaged vesicles for autophagy to defend cells against bacterial invasion. Nature 482: 414–418

Tiwari S, Choi HP, Matsuzawa T, Pypaert M & MacMicking JD (2009) Targeting of the GTPase Irgm1 to the phagosomal membrane via PtdIns(3,4)P2 and PtdIns(3,4,5)P3 promotes immunity to mycobacteria. Nat. Immunol. 10: 907–917

Traver MK, Henry SC, Cantillana V, Oliver T, Hunn JP, Howard JC, Beer S, Pfeffer K, Coers J & Taylor GA (2011) Immunity-related GTPase M (IRGM) Proteins Influence the Localization of Guanylate-binding Protein 2 (GBP2) by Modulating Macroautophagy. J. Biol. Chem. 286: 30471–30480

Tretina K, Park E, Maminska A & Macmicking JD (2019) Interferon-induced guanylate-binding proteins: Guardians of host defense in health and disease. J. Exp. Med.: 1–19

Tripal P, Bauer M, Naschberger E, Mörtinger T, Hohenadl C, Cornali E, Thurau M & Stürzl M (2007) Unique features of different members of the human guanylate-binding protein family. J. Interf. cytokine Res. 27: 44–52

Venegas C, Kumar S, Franklin BS, Dierkes T, Brinkschulte R, Tejera D, Vieira-Saecker A, Schwartz S, Santarelli F, Kummer MP, Griep A, Gelpi E, Beilharz M, Riedel D, Golenbock DT, Geyer M, Walter J, Latz E & Heneka MT (2017) Microglia-derived ASC specks crossseed amyloid-β in Alzheimer’s disease. Nature 552: 355–361

Virreira Winter S, Niedelman W, Jensen KD, Rosowski EE, Julien L, Spooner E, Caradonna K, Burleigh BA, Saeij JPJJ, Ploegh HL & Frickel E-MM (2011) Determinants of GBP Recruitment to Toxoplasma gondii Vacuoles and the Parasitic Factors That Control It. PLoS One 6: e24434

Wallet P, Benaoudia S, Mosnier A, Lagrange B, Martin A, Lindgren H, Golovliov I, Michal F, Basso P, Djebali S, Provost A, Allatif O, Meunier E, Broz P, Py F, Faudry E, Sjo A, Henry T, Yamamoto M & Lyon CB (2017) IFN-γ extends the immune functions of Guanylate Binding Proteins to inflammasome-independent antibacterial activities during Francisella novicida infection. PLoS Pathog.: 1–26

Wandel MP, Pathe C, Werner EI, Ellison CJ, Boyle KB, Malsburg A von der, Rohde J & Randow F (2017) GBPs Inhibit Motility of Shigella flexneri but Are Targeted for Degradation by the Bacterial Ubiquitin Ligase IpaH9.8. Cell Host Microbe 22: 507–518.e5

Wehner M, Kunzelmann S & Herrmann C (2012) The guanine cap of human guanylate-binding protein 1 is responsible for dimerization and self-activation of GTP hydrolysis. FEBS J. 279: 203–210

Yamamoto M, Okuyama M, Ma JS, Kimura T, Kamiyama N, Saiga H, Ohshima J, Sasai M, Kayama H, Okamoto T, Huang DCSS, Soldati-Favre D, Horie K, Takeda J & Takeda K (2012) A cluster of interferon-gamma-inducible p65 gtpases plays a critical role in host defense against toxoplasma gondii. Immunity 37: 302–313

Young J, Dominicus C, Wagener J, Butterworth S, Ye X, Kelly G, Ordan M, Saunders B, Instrell R, Howell M, Stewart A & Treeck M (2019) A CRISPR platform for targeted in vivo screens identifies Toxoplasma gondii virulence factors in mice. Nat. Commun. 10:

Zwack EE, Feeley EM, Burton AR, Baofeng H, Yamamoto M, Kanneganti T-D, Bliska JB, Coers J & Brodsky IE (2017) Guanylate Binding Proteins Regulate Inflammasome Activation in Response to Hyperinjected Yersinia Translocon Components. Infect. Immun. 85: e00778–16

